# New fossils of *Australopithecus sediba* reveal a nearly complete lower back

**DOI:** 10.1101/2021.05.27.445933

**Authors:** Scott A. Williams, Thomas C. Prang, Marc R. Meyer, Thierra K. Nalley, Renier Van Der Merwe, Christopher Yelverton, Daniel García-Martínez, Gabrielle A. Russo, Kelly R. Ostrofsky, Jennifer Eyre, Mark Grabowski, Shahed Nalla, Markus Bastir, Peter Schmid, Steven E. Churchill, Lee R. Berger

**Author notes:** Corresponding author: Scott A. Williams.

## Abstract

Adaptations of the lower back to bipedalism are frequently discussed but infrequently demonstrated in early fossil hominins. Newly discovered lumbar vertebrae contribute to a near-complete lower back of Malapa Hominin 2 (MH2), offering additional insights into posture and locomotion in *Australopithecus sediba*. We show that MH2 demonstrates a lower back consistent with human-like lumbar lordosis and other adaptations to bipedalism, including an increase in the width of intervertebral articular facets from the upper to lower lumbar column (“pyramidal configuration”). This contrasts with recent work on lordosis in fossil hominins, where MH2 was argued to demonstrate no appreciable lordosis (“hypolordosis”) similar to Neandertals. Our three-dimensional geometric morphometric (3D GM) analyses show that MH2’s nearly complete middle lumbar vertebra is human-like in shape but bears large, cranially-directed transverse processes, implying powerful trunk musculature. We interpret this combination of features to indicate that *A. sediba* used its lower back in both human-like bipedalism and ape-like arboreal positional behaviors, as previously suggested based on multiple lines of evidence from other parts of the skeleton and reconstructed paleobiology of *A. sediba*.

## 1. Introduction

Bipedal locomotion is thought to be one of the earliest and most extensive adaptations in the hominin lineage, potentially evolving initially 6-7 million years (Ma) ago. Bipedalism evolved gradually, however, and hominins appear to have been facultative bipeds on the ground and competent tree climbers for at least the first 4.4 Ma of their existence(1, 2). How long climbing adaptations persisted in hominins and when obligate bipedalism evolved are major outstanding questions in paleoanthropology. *Australopithecus sediba—*a 2 Ma australopith from the site of Malapa, Gauteng province, South Africa—has featured prominently in these discussions, as well as those concerning the origins of the genus *Homo(3)*.

Previous studies support the hypothesis that *A. sediba*, while clearly bipedal, possessed adaptations to arboreal locomotion and lacked traits reflecting a form of obligate terrestriality observed in later hominins(4–8). Malapa Hominin 2 (MH2) metacarpals are characterized by relative trabecular density most similar to orangutans, which suggests power grasping capabilities(9), and the MH2 ulna was estimated to reflect a high proportion of forelimb suspension in the locomotor repertoire of *A. sediba*(10). Evidence from the lower limb also suggests that *A. sediba* lacked a robust calcaneal tuber(5) and a longitudinal arch(6), both thought to be adaptations to obligate, human-like bipedalism, and demonstrates evidence for a mobile subtalar joint proposed to be adaptively significant for vertical climbing and other arboreal locomotor behaviors(7, 11–13). The upper thorax(4), scapula(14, 15), and cervical vertebrae(16) of *A. sediba* suggest shoulder and arm elevation indicative of arboreal positional behaviors. Furthermore, analysis of dental calculus from Malapa Hominin 1 (MH1) indicates that this individual’s diet was high in C_3_ plants like fruit and leaves, similar to savannah chimpanzees and *Ardipithecus ramidus*(17).

Despite the presence of climbing adaptations, *A. sediba* demonstrates clear evidence for bipedal locomotion. These adaptations are present not just in the lower limb(11–13), but also in the lower back, including strong dorsal (lordotic) wedging of the two lower lumbar vertebrae, which contribute to a lordotic (ventrally curved) lower back(18, 19). However, the initial recovery of just the last two lumbar vertebrae of MH2 limited interpretations of spinal curvature, and a study of the MH2 pelvis reconstruction(20) suggested that *A. sediba* was characterized by a small lordosis angle estimated from calculated pelvic incidence(21). The fragmentary preservation of previously known lumbar vertebrae prevented comparative analysis of overall lower back morphology and allowed limited interpretations of *A. sediba* positional behavior.

Here, we report the discovery of portions of four lumbar vertebrae from two *ex situ* breccia blocks that were excavated from an early 20^th^ century mining road and dump at Malapa. The mining road (approximately three meters in width) is located in the northern section of the site two meters north of the main pit that yielded the original *A. sediba* finds(22). The road transverses the site in an east-west direction and was constructed using breccia and soil removed from the main pit by the historic limestone miners. Specimens U.W.88-232, -233, and -234 were recovered in 2015 from the upper section of layer 2 (at a depth of 10 cm) and formed part of the foundation layer of the road. The road can be distinguished from the surrounding deposits by a section of compacted soil (comprising quartz, cherts, and flowstone) and breccia that extends between layers 1 and 2. Breccia recovered from the road, including the block containing U.W.88-232, 233, and -234 similarly presented with quantities of embedded quartz fragments and grains. The breccia block containing specimens U.W.88-280 and U.W.88-281, along with U.W.88-43,-44, and -114 (Williams et al., 2013, 2018), were recovered from the miner’s dump comprised of excess material (soil and breccia) used for the construction of the miner’s road. The composition of the road matrix and associated breccia, as well as the breccia initially recovered from the mine dump, corresponds to the facies D and E identified in the main pit(22). Facies D includes a fossil-rich breccia deposit that contained the fossil material associated with MH2(22). Therefore, the geological evidence suggests that the material used for the construction of the miner’s road was sourced on-site, and most probably originated from the northern section of the main pit.

The new vertebrae (second and third lumbar) are preserved in articulation with each other (Fig. 1) and refit at multiple contacts with the previously known penultimate (fourth) lumbar vertebra (Fig. 2). Together, the new and previously known(18, 19) vertebral elements form a continuous series from the antepenultimate thoracic vertebra through the fifth sacral element, with only the first lumbar vertebra missing major components of morphology (Figs. 2–3; Supplementary Fig. 1). The presence of a nearly complete lower back of a female *A. sediba* individual allows us to test existing hypotheses about the functional morphology and evolution of purported adaptations to bipedalism in fossil hominins, including overall lumbar vertebra shape, lumbar lordosis, and progressive widening of the articular facets and laminae (pyramidal configuration) of the lower back.

**Figure 1.**
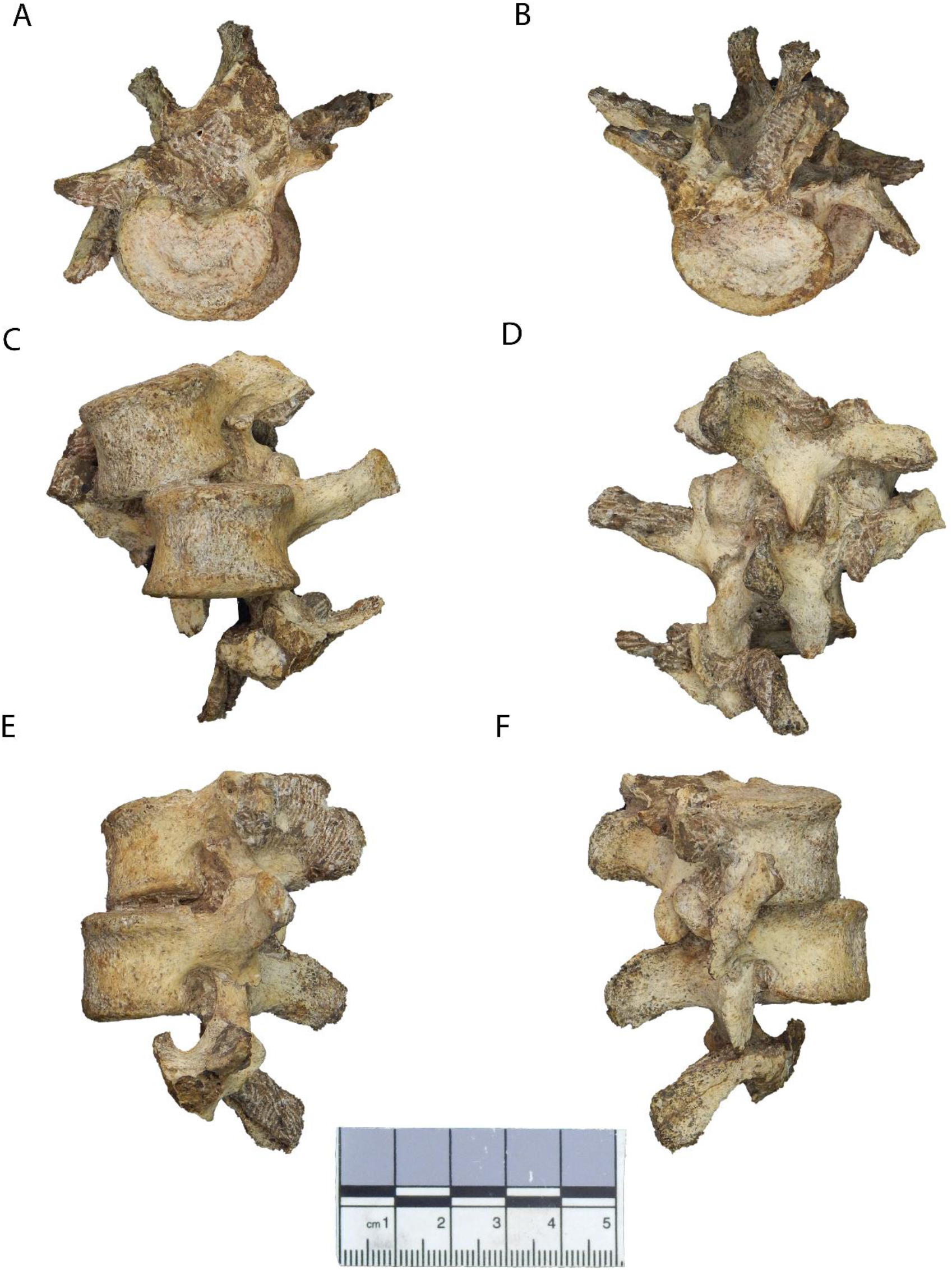
New lumbar vertebrae of Malapa Hominin 2 (MH2). Vertebrae in **(A)** superior, **(B)** inferior, **(C)** ventral, **(D)** dorsal, **(E)** left lateral, and **(F)** right lateral views. The partial inferior articular facets of the first lumbar vertebra are embedded in matrix (see Supplementary Fig. 2). The second lumbar vertebra (U.W.88-232) is in the superior-most (top) position, the third lumbar vertebra (U.W.88-233) is in the middle, and portions of the upper neural arch of the fourth lumbar vertebra (U.W.88-234) are in the inferior-most (bottom) position. The lumbar block is rotated to center U.W.88-233 in all views. These fossils are curated and available for study at the University of the Witwatersrand.

**Figure 2.**
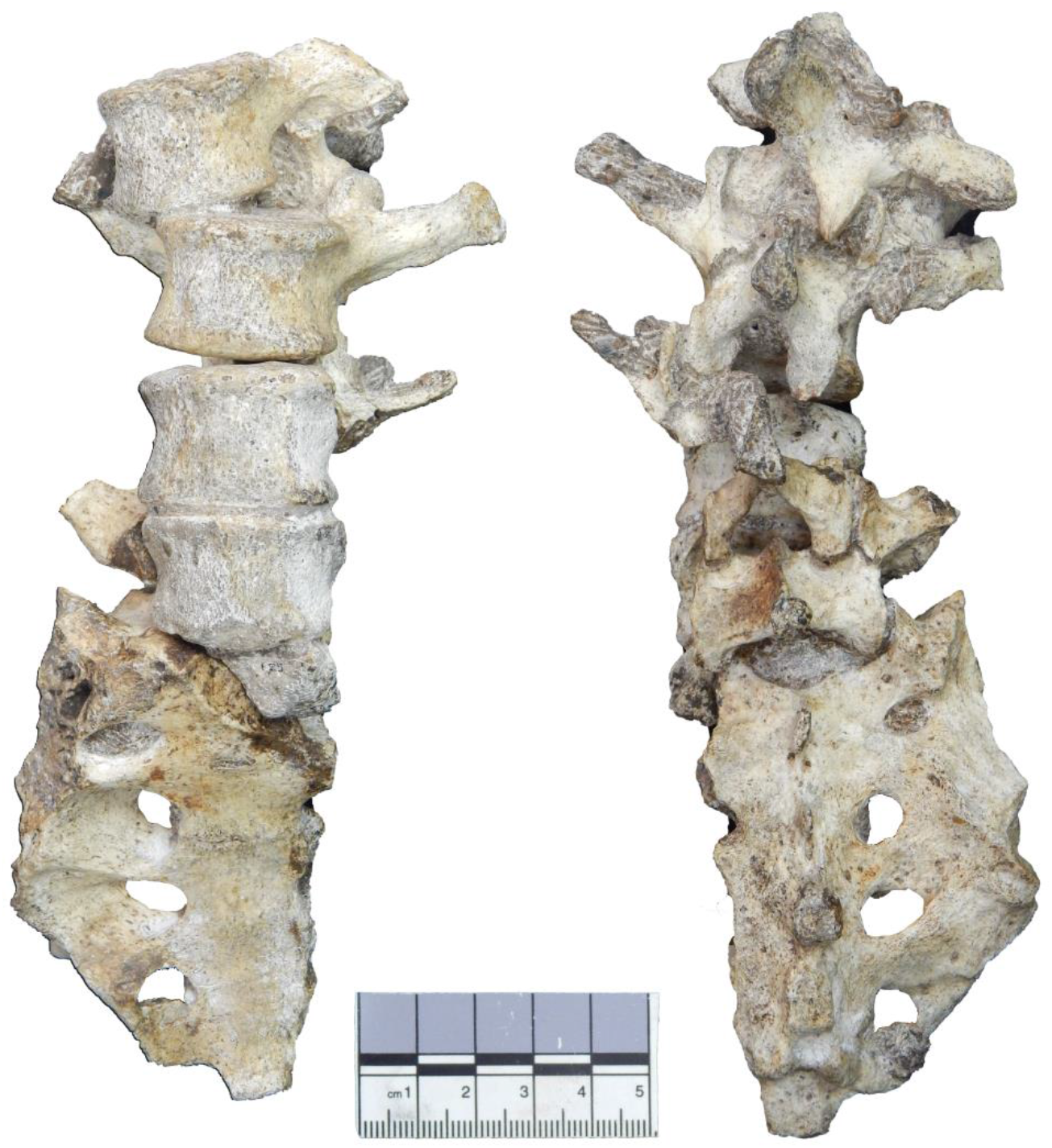
The lower back of Malapa Hominin 2 in ventral (left) and dorsal (right) views. New second and third lumbar vertebrae (U.W.88-232, U.W.88-233) are positioned at the top, and U.W.88-234 contributes to the upper portion of the fourth lumbar vertebra (U.W.88-127/153/234). The fifth lumbar vertebra (U.W.88-126/138) sits atop the sacrum (U.W.88-137/125). The lower back elements are preserved together in three blocks, each containing multiple elements held together in matrix and/or in partial articulation: 1) most of the sacrum (U.W.88-137), the neural arch of L5 (U.W.88-153), the inferior portion of the neural arch of L4 (U.W.88-138); 2) the L4 (U.W.88-127) and L5 (U.W.88-126) vertebral bodies, and partial S1 body (U.W.88-125); 3) L1 inferior neural arch (U.W.88-281; concealed in matrix), L2 (U.W.88-232), L3 (U.W.88-233), and upper neural arch of L4 (U.W.88-234). The vertebral body fragment of L1 (U.W.88-280) is preserved within the matrix of a separate block containing the lower thoracic vertebrae (U.W.88-43/114 and U.W.88-44) (See Fig. 3; Supplementary Figs. 1–2).

**Figure 3.**
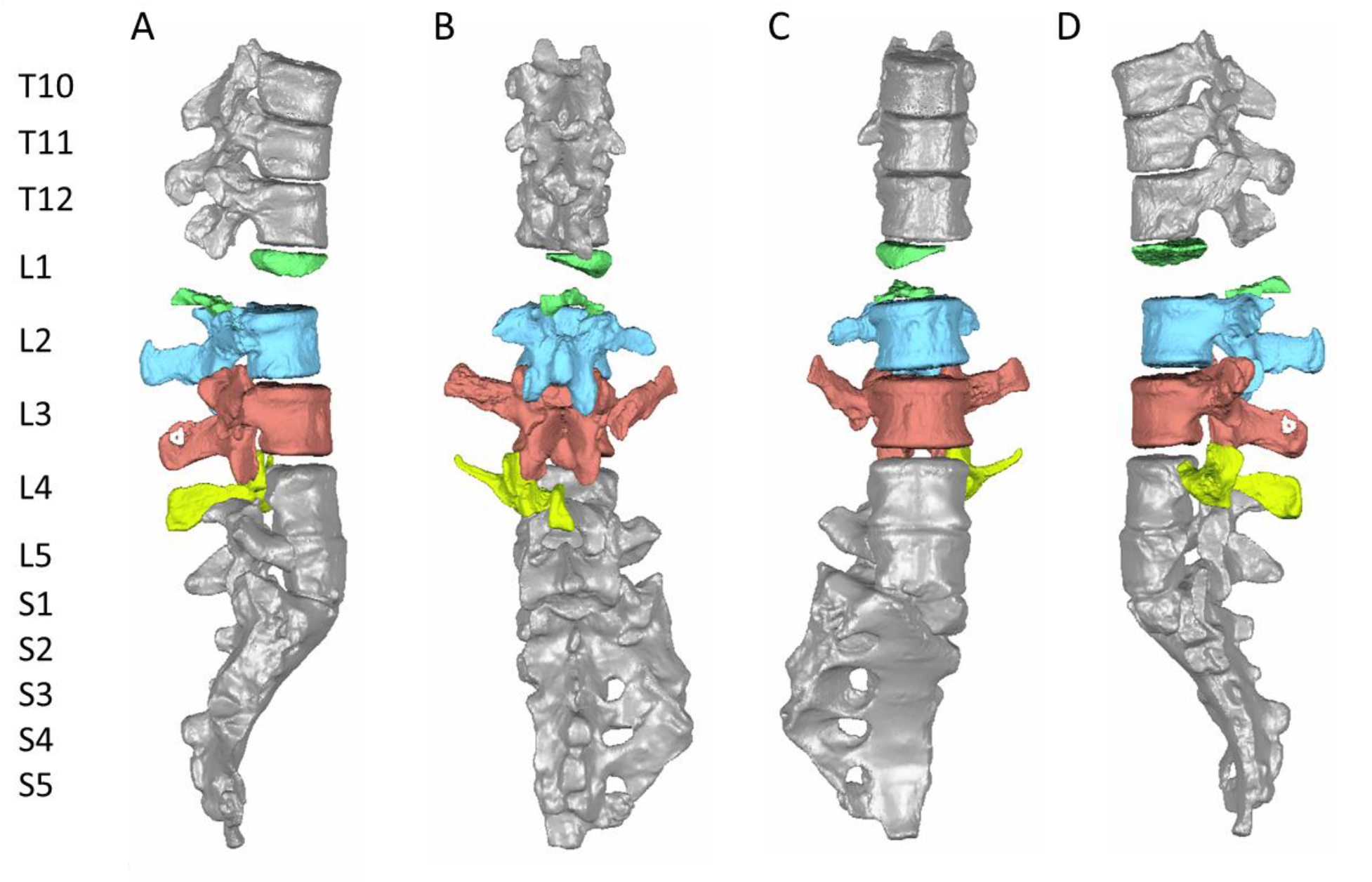
Surface renderings of the new lumbar vertebrae positioned between previously known vertebrae. Components of the L1 vertebra (U.W.88-280/281) are shown in green, the L2 (U.W.88-232) is shown in blue, the L3 (U.W.88-233) is shown in red, and the new component of L4 (U.W.88-234) is shown in yellow. The previously known aspects of L4 (U.W.88-127/153), the L5 (U.W.88-126/138), sacrum (U.W.88-137/125), and lower thoracic vertebrae (T10-T12) are fully published in Williams et al. (2018). Fossils are shown in their original condition and refitted at contact points but not reconstructed. Three-dimensional models of all these vertebrae can be downloaded on MorphoSource.org.

## Results

The five new fossils, U.W.88-232, -233, -234, -280, and -281, are described below and shown in Figs. 1–3 and Supplementary Figs. 1–2. Measurements are included in Table 1. A depiction of the anatomical features mentioned in the descriptions below and throughout the manuscript is shown in Fig. 4).

**Table 1.** Measurements (in mm for linear measurements and degrees for angles; measurement definitions are included in the Supplementary Material) on lumbar vertebrae.

|  | U.W. 88-232<br>(L2) | U.W. 88-233<br>(L3) | U.W. 88-127/153/<br>234<br>(L4) | U.W. 88-126/138<br>(L5) |
| --- | --- | --- | --- | --- |
| 1. Body sup. transv. width (MαW7) | 29.5 | 30.1 | 31.4 | 32.8 |
| 2. Body sup. dorsoven. dia. (Mα4) | 20.8 | 21.4 | 22.2 | 21.4 |
| 3. Body inf. transv. width (Mα8) | 29.0 | 31.4 | 32.4 | 28.8 |
| 4. Body inf. dorsoven. dia. (MαW5) | 21.1 | 21.0 | 21.2 | 19.8 |
| 5. Body ventral height (MαW1) | 21.0 | 21.75 | 22.1 | 21.0 |
| 6. Body dorsal height (MαW2) | 22.5 | 22.25 | 21.5 | 17.0 |
| 7. Body wedging angle (in degrees) | 4.1° | 1.3° | -1.6° | -10.7° |
| 8. Spinal canal dorsoven. dia. (MαW10) | 10.5 | 8.85 | - | 23 |
| 9. Spinal canal transv. dia. (MαW11) | 17.6 | 17.3 | - | 16.3 |
| 10. Sup.-inf. inter-AF height | - | 37.0 | 32.6 | 31.5 |
| 11. Max. inter-SAF dist. | - | 24.0 | - | 28.5 |
| 12. Min. inter-SAF dist. | - | 14.5 | - | - |
| 13. Max. inter-IAF dist. | 23.0 | 25.0 | (28.0) | (33.0) |
| 14. Min. inter-IAF dist. | 11.0 | 9.5 | 11.6 | 15.6 |
| 15. SAF sup.-inf. dia. | - | 12.8 | - | 13.4 |
| 16. SAF transv. dia. | - | 11.5 | - | 10.8 |
| 17. IAF sup.-inf. dia. | 11.5 | 11.5 | 14.7 | 14.4 |
| 18. IAF transv. dia. | 8.1 | 8.9 | 9.2 | 11.7 |
| 19. Spinous process angle (MαW12) | 176° | 160° | 163° | 166° |
| 20. Spinous process length (MαW13) | 27.0 | 28.0 | 28.0 | 23.6 |
| 21. Spinous process terminal trans. width | 6.9 | 7.4 | 8.1 | 6.85 |
| 22. Spinous process terminal sup.-inf. height | 13.8 | 11.75 | 12.7 | 7.15 |
| 23. Transverse process base sup.-inf. height | 11.5 | 12.2 | - | 13.9 |
| 24. Transverse process angle | 78° | 82.0° | - | 50° |
| 25. Transverse process length | - | 31.0 | - | - |
| 26. SAF orientation (in degrees) | - | 31° | 33° | 26° |
| 27. Pedicle sup.-inf. height | 10.9 | 10.6 | - | 11.2 |
| 28. Pedicle transv. width | 5.9 | 7.1 | 9.0 | 10.9 |
| 29. Pedicle dorsoven. length | 5.0 | 5.6 | 6.5 | 7 |
| 30. Lamina sup.-inf. height | 16.1 | 15.4 | - | 14 |
| 31. Lamina transv. width | 20.0 | 22.0 | - | 30.5 |

<sup>1</sup> Parentheses indicate that the structure is incomplete and its measurement is estimated.

**Figure 4.**
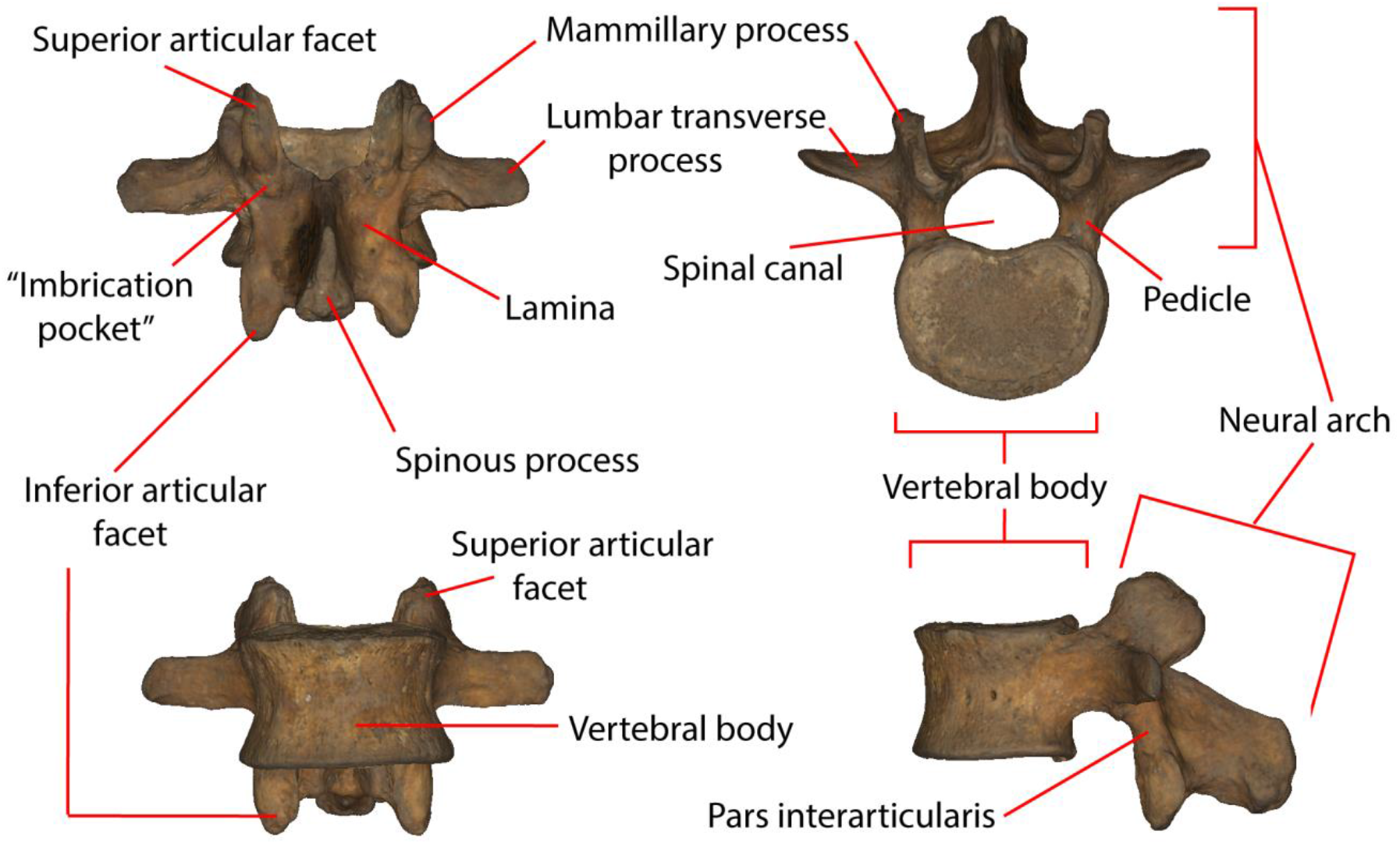
Anatomical features of lumbar vertebrae. A human middle lumbar vertebra is shown in dorsal (upper left), cranial (upper right), ventral (lower left), and lateral (lower right) views.

### Descriptions of new fossil material

U.W.88-280: This is a partial, superior portion of a vertebral body concealed in the matrix of the lower thoracic block, which also contains U.W.88-114, -43, and -44 (antepenultimate, penultimate, and ultimate thoracic vertebrae). U.W.88-280 was revealed in the segmentation of micro-CT (hereafter, mCT) data. U.W.88-280 represents the right side of an upper vertebral body, which we identify as part of the first lumbar vertebra of MH2, and its preservation approaches the sagittal midline. The preserved portions measure 16.5 mm dorsoventrally and 14.0 mm mediolaterally at their maximum lengths. The lateral portion of the vertebral body is only preserved ~5.0 mm inferiorly from the superior surface, but there is no indication of a costal facet on the preserved portion.

U.W.88-281: This is the partial neural arch of a post-transitional, upper lumbar vertebra concealed in matrix at the top of the new lumbar block. It was revealed through the segmentation of mCT data. It consists of the base and caudal portion of the spinous process and parts of the inferior articular processes. The remainder of the vertebra is sheared off and unaccounted for in the new lumbar block. U.W.88-281 is fixed in partial articulation with the subjacent second lumbar vertebra (L2), U.W.88-232. Therefore, we identify U.W.88-281 as part of the first lumbar vertebra based on its morphology and position within the block. The left inferior articular facet is more complete than the right, with approximately 6.0 mm of its superior-inferior (SI) height preserved, and is complete mediolaterally, measuring ~8.0 mm in width. The minimum distance between the inferior articular facets is 12.5 mm, and the maximum preserved distance between them is 21.75 mm. The preserved portion of the spinous process is 12.75 mm in dorsoventral length.

U.W.88-232: This is the L2 discovered in the mining road block. It remains in articulation with the third lumbar vertebra (L3), U.W.88-233, held together with matrix. Some portions of U.W.88-232 are covered by adhering matrix or other fossil elements (U.W.88-281 and -282), so mCT data were used to visualize the whole vertebra (Supplementary Fig. 1). U.W.88-232 is mostly complete, missing the cranial portions of its superior articular processes and distal portions of its lumbar transverse processes. It is distorted due to crushing dorsally from the right side and related breakage and slight displacements of the left superior articular process at the *pars interarticularis* and the right lumbar transverse process at its base. Although broken at its base and displaced slightly ventrally, the right lumbar transverse process is more complete than the left side, which is broken and missing ~10 mm from its base. As a consequence of crushing, the neural arch is displaced towards the left side, and the spinal canal is significantly distorted. A partial mammillary process is present on the left superior articular process, sheared off along with the remainder of the right superior articular process ~8 mm from its base. The left side is similar but much of the mammillary process is sheared off in the same plane as the right side, leaving only its base on the lateral aspect of the right superior articular process. The vertebral body is complete and undistorted, and the spinous process and inferior articular processes are likewise complete but affected by distortion. Standard measurements of undistorted morphologies are reported in Table 1.

U.W.88-233: This is the L3 and the most complete vertebra in the lumbar series, although some aspects of the neural arch are distorted, broken, and displaced. It is held in matrix and partial articulation with U.W.88-234, the subjacent partial fourth lumbar vertebra (L4). Due to its position between articulated elements U.W.88-232 and -234 and some adhering matrix, U.W.88-233 was visualized using mCT data. U.W.88-233 is essentially complete; however, like U.W.88-232, the neural arch is crushed from the dorsal direction, with breaks and displacement across the right *pars interarticularis* and the right lumbar transverse process at is base, with additional buckling in the area of the latter near the base of the of the right superior articular process, resulting in a crushing of the spinal canal. The vertebral body, pedicles, spinous process, and superior and inferior articular processes are complete, as are the lamina and lumbar transverse processes aside from the aforementioned breakage. The left lumbar transverse process is unaffected by taphonomic distortion. Standard measurements of undistorted morphologies are reported in Table 1.

U.W.88-234: This is a partial neural arch of the previously known penultimate lumbar vertebra (U.W.88-127/153). U.W.88-234 refits in two places with the previously known L4 vertebra, its partial pedicle with the vertebral body (U.W.88-127) and its spinous process with the inferior base of the spinous process and inferior articular processes (U.W.88-153) (Figs. 2–3). Only the spinous process and right pedicle, lumbar transverse process, superior articular processes, and partial lamina are present and in articulation with U.W.88-233. Matrix adheres to the spinous process and lumbar transverse process, so for this element mCT data were used to visualize and virtually refit it with U.W.88-127/153, forming a partial L4 missing the left superior articular process, lumbar transverse process, most of the pedicle, the right lateral aspect of the inferior articular process, a portion of the lamina, the inferior aspect of the lumbar transverse process, and a wedge shaped area of the lateral body-pedicle border. Preserved standard measurements are reported in Table 1.

### Middle lumbar vertebra comparative morphology

The new middle lumbar vertebra, U.W.88-233, is complete, and although the neural arch is compressed ventrally into the spinal canal space, it can be reasonably reconstructed from mCT data (see Methods). We used three-dimensional geometric morphometrics (3D GM) to evaluate the shape affinities of U.W.88-233 among humans, great apes, and fossil hominins. The results of our principal components analysis (PCA) on Procrustes-aligned shape coordinates reveal that *A. sediba* falls within or near the human distribution on the first three principal components (PC 1-3) (Fig. 5). PC1 explains 31% of the variance in the dataset, and along it hominins are characterized by more sagittally oriented and concave superior articular facets, more dorsally oriented transverse processes, a dorsoventrally shorter and cranially oriented spinous process, a slightly larger and craniocaudally shorter vertebral body, and more caudally positioned superior and inferior articular facets relative to the vertebral body compared to great apes.

**Figure 5.**
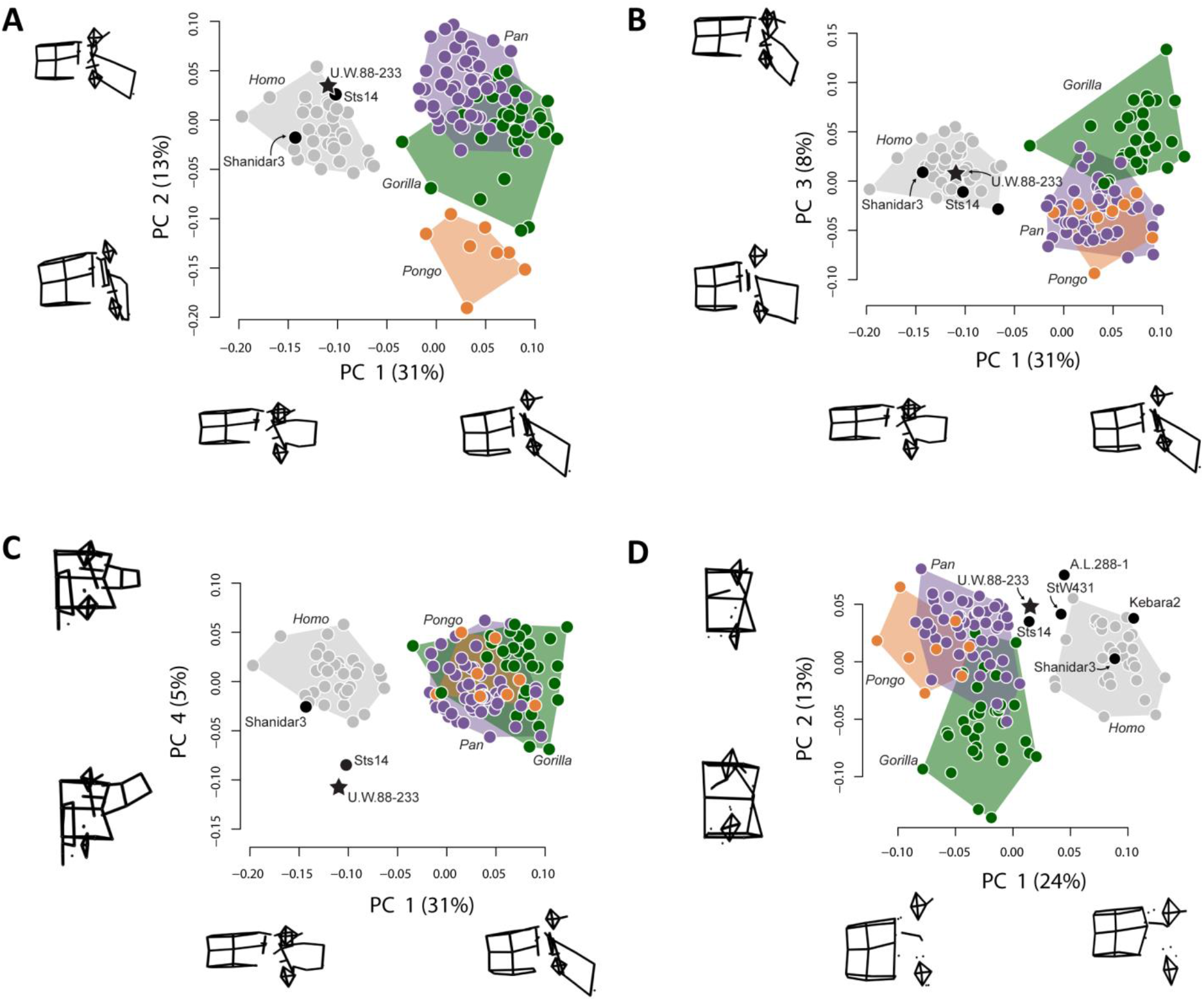
Principal components analysis on middle lumbar vertebra 3D landmark data. **(A-C)** PCA on the full set of 48 landmarks, including Sts 14 (*A. africanus*), U.W.88-233 (*A. sediba*), and Shanidar 3 (Neandertal). **(A-B)** Hominins separate from great apes on PC1 (wireframes in lateral view), African apes and hominins separate from orangutans on PC2 (wireframes in lateral view), and **(C)** australopiths separate from other hominids on PC4 (wireframes in posterior view). Note that spinous and transverse process lengths and orientations drive much of the variance in middle lumbar vertebrae. **(D)** PCA on a reduced landmark set (sans and spinous and transverse process landmarks) to include A.L. 288-1 (*A. afarensis*), StW 431 (*A. africanus*), and Kebara 2 (Neandertal). Notice that australopiths fall outside the modern human convex hulls, with Sts 14 and MH2 close to those of the African apes. 3D landmark data were subjected to Procrustes transformation.

PC2 explains 13% of the variance and contrasts long spinous processes and relatively neutrally wedged vertebral bodies of hominins and African apes with the shorter spinous processes and strongly ventrally wedged vertebral bodies of orangutans. PC3 explains 8% of the variance and largely contrasts robust vertebral bodies with caudally oriented spinous processes in gorillas with more gracile vertebral bodies and less caudally oriented spinous processes in chimpanzees and orangutans; hominins fall intermediate between these groups. PC4 explains 5% of the variance, and contrasts *A. sediba* and *A. africanus* with both humans and great apes. Sts 14 and especially U.W.88-233 are characterized by longer, taller, more cranially oriented lumbar transverse processes that do not taper distally and more sagittally oriented (as opposed to more coronally oriented) articular facets (Fig. 5; Supplementary Fig. 3). Therefore, the vertebral body and spinous process of U.W.88-233 are human-like in overall shape, but the lumbar transverse processes are robust, cranially oriented, and cranially positioned on the pedicles.

To include other fossil hominins with broken processes, we ran a second 3D GM analysis excluding the majority of lumbar transverse process and spinous process landmarks. This analysis, which includes landmarks on the vertebral body, superior and inferior articular facets, and the bases of the transverse and spinous processes, produces a similar pattern compared to the analysis on the full landmark set (Fig. 5). Humans and great apes separate along PC1, which is largely explained by vertebral body height and SI position of the articular facets relative to the vertebral body. U.W.88-233 and Sts 14 fall intermediately between modern humans and great apes along PC1.

We used a Procrustes distance-based Analysis of Variance (ANOVA) to evaluate the effect of centroid size on lumbar shape(23). The results show significant effects of centroid size (F = 9.83; *p* < 0.001), genus (with hominins pooled; F = 27.7; *p* < 0.001), and an interaction between genus and centroid size (F = 1.48; *p* = 0.01), implying unique shape allometries within genera (Table 2). We plotted standardized shape scores derived from a multivariate regression of shape on centroid size against centroid size to visualize shape changes(24) (Supplementary Fig. 4). In general, larger centroid sizes are associated with 3D shape changes including dorsoventrally longer and more caudally projecting spinous processes, more cranially oriented and less sagittally oriented lumbar transverse processes, and less caudally projecting inferior articular facets. Since body mass scales as the cube of linear dimensions and the physiological cross-sectional area of skeletal muscle—a major determinant of isometric force production—scales as the square of linear dimensions, larger individuals should be relatively weaker with all else held equal. The length increases of the spinous process and lumbar transverse processes act to increase the moment arms of the erector spinae and quadratus lumborum muscles, respectively, resulting in greater moments that contribute to lumbar extension, ventral flexion, and lateral flexion to cope with increases in body mass. Importantly, however, the cranially oriented lumbar transverse processes of U.W.88-233 (and Sts 14) appear not to be explained by centroid size given its relatively small size and overlap with *Pan* in standardized shape scores (Supplementary Fig. 4).

**Table 2.**
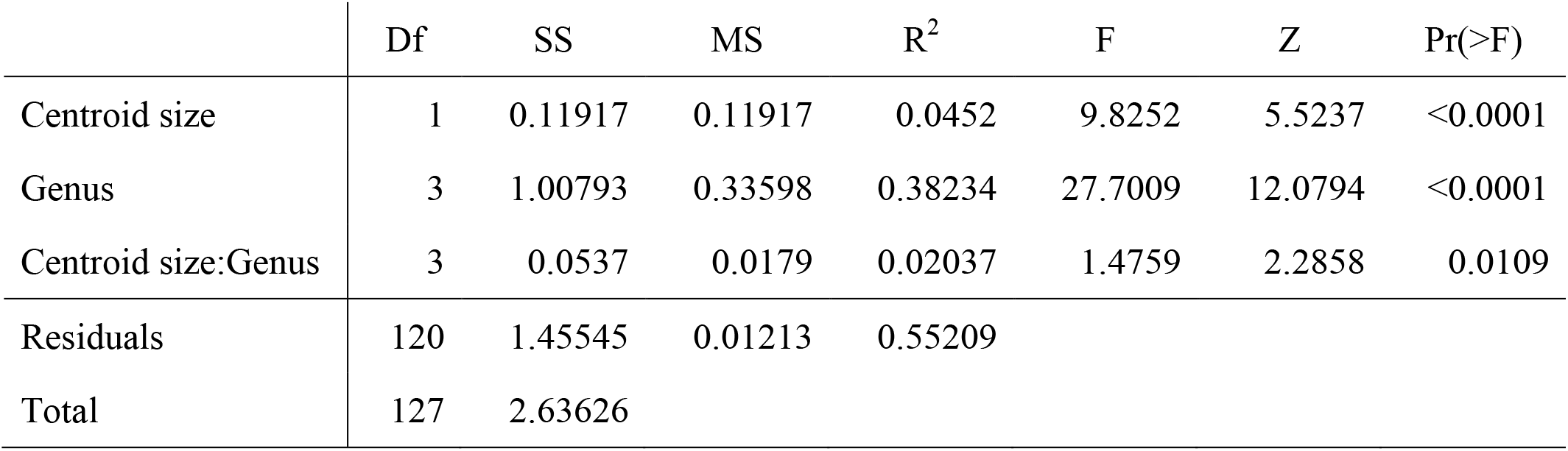
| Procrustes ANOVA results of centroid size and middle lumbar vertebra shape.

### Wedging angles and inferred lumbar lordosis

Wedging of vertebrae contribute to multiple sagittal curvatures of the human spine, with dorsal wedging of lower lumbar vertebrae contributing to a ventral curvature of the lumbar spine (lumbar lordosis). This sinusoidal configuration passively balances the upper body over the pelvis and allows for the unique system of weight bearing and force transmission found in members of the human lineage (25–31). Wedging angles for individual lumbar vertebrae (L2-L5) and sum L2-L5 wedging were calculated for *A. sediba* and the comparative sample. MH2 produces the greatest sum wedging value of any adult early hominin (−6.8°) and falls within the 95% confidence intervals of female modern humans (Fig. 6). Sexual dimorphism is observed in *A. africanus* (31), with inferred female Sts 14 falling within the 95% confidence intervals of female modern humans and inferred male *A. africanus* StW 431 (but see refs. (32, 33)) falling within the 95% confidence intervals of modern human males (Fig. 6). Two male Neandertal lumbar series demonstrate strong kyphotic wedging and fall outside the modern human confidence intervals and within or near those of chimpanzees (see Fig. 6 caption).

**Figure 6.**
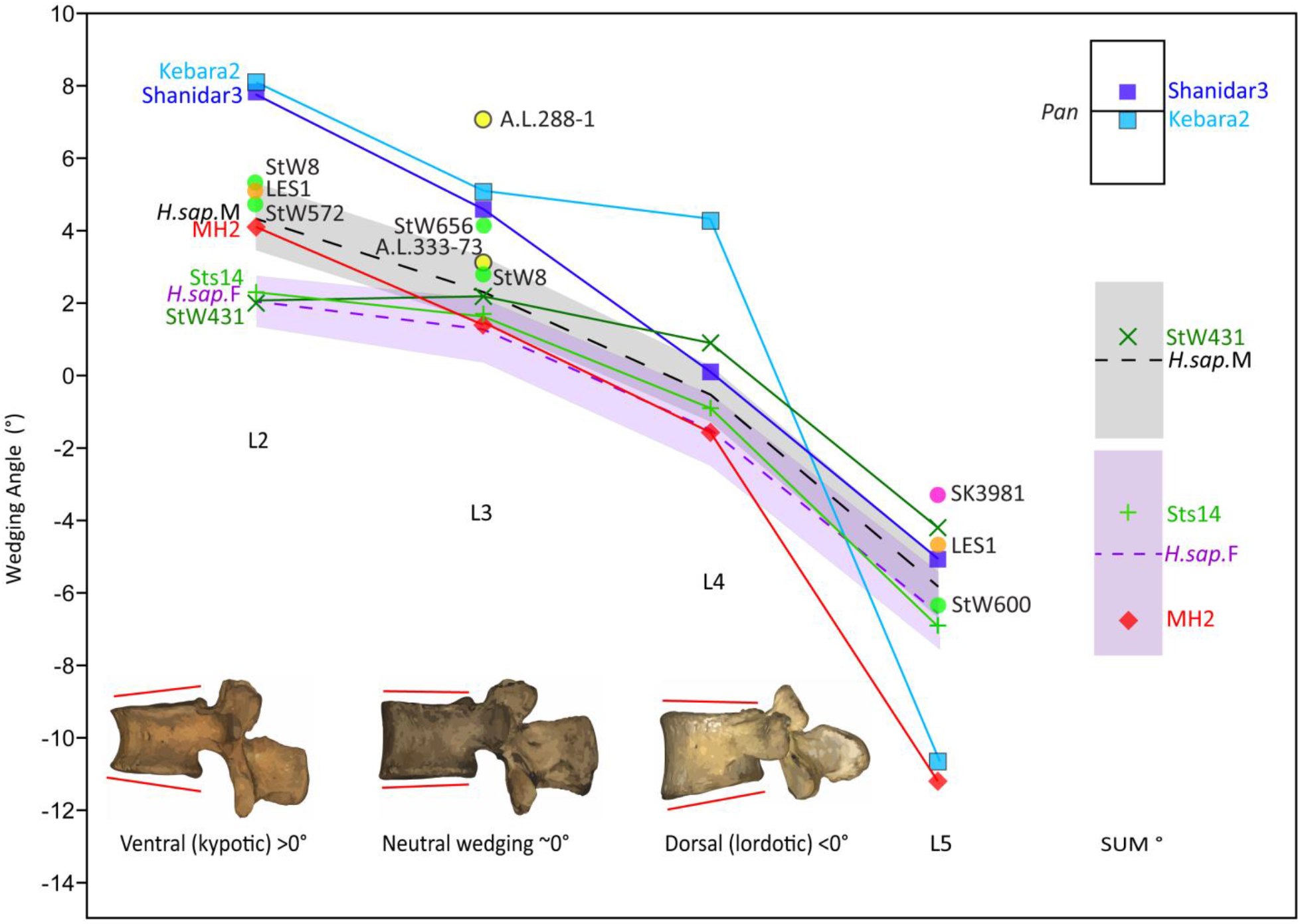
Lumbar vertebral body wedging angles. Vertebral wedging angles are plotted from L2 through L5. Specimens with a full complement of L2-L5 vertebrae are labelled on the left side (*A. africanus*: Sts 14, StW 431; *A. sediba*: MH2; Neandertals: Kebara 2, Shanidar 3). Fossil individuals represented by one to three L2-L5 vertebrae are labelled at each vertebral level (*A. afarensis*: A.L. 288-1, A.L. 333-73; *A. africanus*: StW 8, StW 572, StW 656, StW 600(53); *P. robustus*: SK 3981; *H. naledi*: LES1(54)). Human females [L2: *μ*=2.06, *σ*=2.08 (range=−2.07, 5.98); L3: *μ*=1.26, *σ*=2.56 (−3.39, 6.59); L4: *μ*=−1.49, *σ*=2.92 (−7.91, 4.63); L5: *μ*=−6.473, *σ*=3.12 (−12.35, 2.21); Sum L2-L5: *μ*=−4.93, *σ*=8.09 (−18.20, 11.19)] and males [L2: *μ*=4.41, *σ*=3.31 (range=−3.39, 12.80); L3: *μ*=2.38, *σ*=−4.83 (−4.83, 11.24); L4: *μ*=−0.52, *σ*=2.76 (−8.25, 4.84); L5: *μ*=−5.86, *σ*=2.91 (−11.54, 1.75); Sum L2-L5: *μ*=0.41, *σ*=7.81 (−15.59, 15.53)] are represented with means and 95% confidence intervals. Only specimens with a full complement of L2-L5 vertebrae are included in the sum wedging angle column. Chimpanzees (*Pan troglodytes*) and gorillas (*Gorilla gorilla*) with four lumbar vertebrae are also included in the sum wedging angle column for comparative purposes.

Patterns of change across lumbar levels demonstrate that MH2’s vertebrae transition from ventral (kyphotic) wedging at the L2 (most similar to male modern humans) and L3 (most similar to female modern humans) levels to dorsal (lordotic) wedging at L4. At the L4 level MH2 is most similar to female modern humans and female *A. africanus* (Sts 14), in contrast with male *A. africanus* (StW 431) and male Neandertals, which reach dorsal wedging only at the last lumbar level (Fig. 6). As shown previously(18), the last lumbar vertebra of MH2 is strongly dorsally wedged like that of the Kebara 2 Neandertal, whereas other fossil hominins do not demonstrate this pattern. MH2 produces a more modern humanlike sum of wedging angles, however, because it falls near the human means levels L2-L4, whereas Neandertals fall well above the human 95% confidence intervals.

### Pyramidal configuration of the neural arch

The recovery of new lumbar vertebrae of MH2 allows for the quantification and comparison of australopith inter-articular facet width increase. Humans are characterized by a pyramidal configuration of the articular facets such that those of lower lumbar vertebrae are progressively transversely wider than those of upper lumbar vertebrae(28). Using an index of the last lumbar-sacrum inter-articular maximum distance relative to that of lumbar vertebrae three levels higher (L2-L3 in hominins, L1-L2 in chimpanzees and gorillas), we show that *A. africanus* (Sts 14 and StW 431; average = 1.42) and *A. sediba* (1.43) fall at the low end of the range of modern human variation in this trait (Fig. 7). Note that A.L. 288-1 (*A. afarensis*) falls at the low end of human variation near other australopiths if the preserved lumbar vertebra (A.L. 288-1aa/ak/al) is treated as an L3(28, 30, 34, 35), but outside the range of human variation and within that of orangutans if it is treated as an L2(36). *H. erectus* and Neandertals fall well within the range of modern human variation.

**Figure 7.**
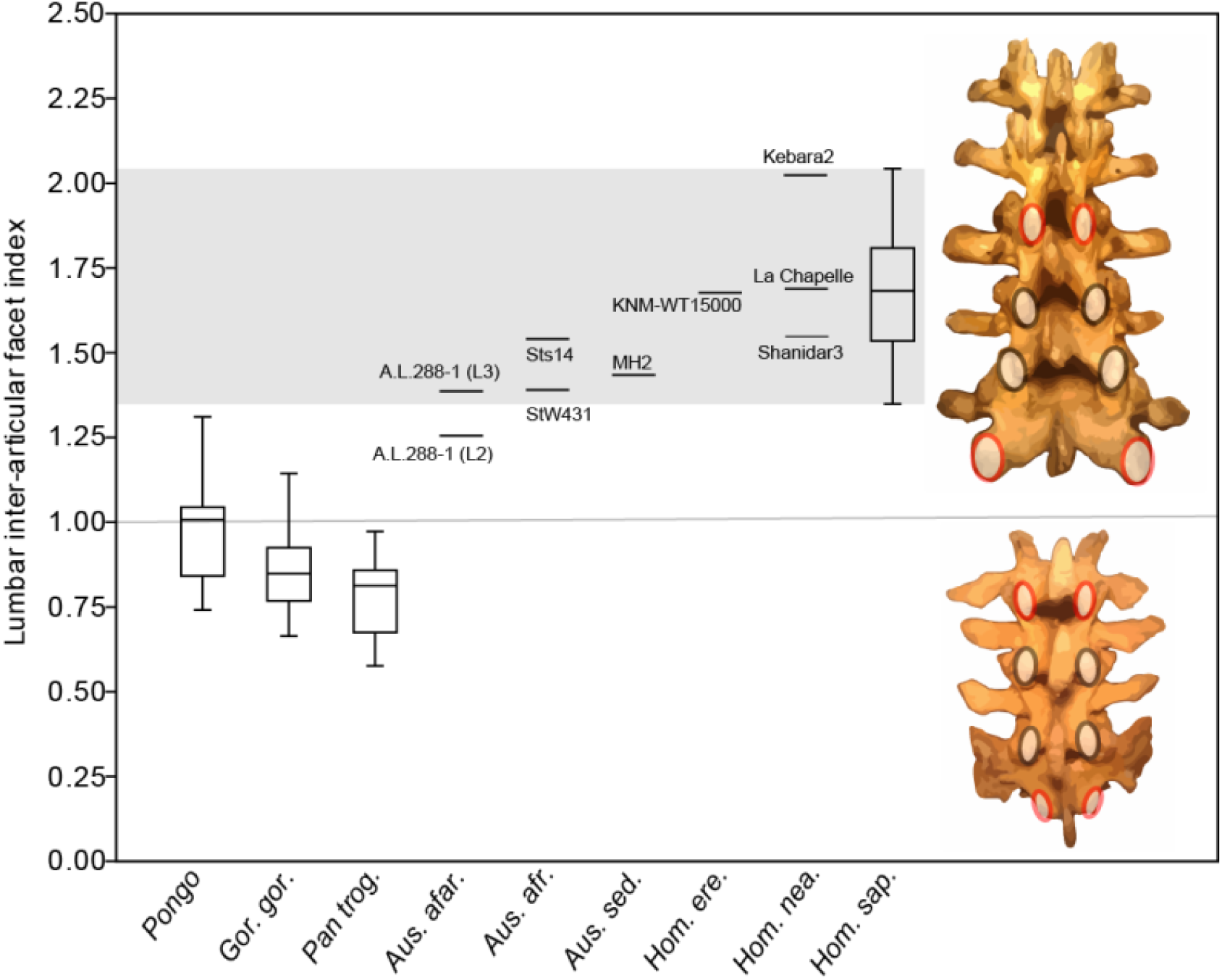
Pyramidal configuration of articular facet spacing in hominids. The inter-articular facets of the last lumbar/sacrum and those of lumbar vertebrae three elements higher in the column (L1-L2 in chimpanzees and gorillas with four lumbar vertebrae; L2-L3 in hominins) are included as the numerator and denominator, respectively, in a lumbar inter-articular facet index. These levels are highlighted in red in both a human (top) and a chimpanzee (bottom). The grey box highlights the range of variation observed in the modern human sample.

## Discussion

The recovery of two new lumbar vertebrae and portions of other vertebrae of adult female *Australopithecus sediba* (MH2), together with previously known vertebrae, form a nearly complete lumbar column (Figs. 2–3) and allows us to test hypotheses based on more limited material. As we outline below, *A. sediba* demonstrates primary adaptations to bipedal locomotion in its lower back, including human-like overall lumbar vertebra shape, evidence for lumbar lordosis in the pattern of bony wedging of lumbar vertebral bodies, and progressive widening of neural arch structures moving caudally (“pyramidal configuration”) of lumbar vertebrae and the sacrum.

The overall morphologies of lumbar vertebrae are informative with regard to locomotion and posture in primates (37–41). Hominoids are characterized by derived vertebral morphologies related to orthogrady and antipronograde positional behaviors, and early hominins have been found to largely resemble modern humans in lumbar vertebra shape, with some retained primitive morphologies (26, 42, 43). Our 3D GM results show that the middle lumbar vertebra of *sediba* (U.W.88-233) falls with humans to the exclusion of great apes in overall shape, but bears large, cranially oriented lumbar transverse processes (Fig. 5). Large lumbar transverse processes provide ample surface area for powerful trunk musculature (26, 42, 44) and in their cranial orientation increase the moment arms and torque generation capabilities of psoas major and quadratus lumborum, and via the middle lumbar fascia, the internal and external obliques and transversus abdominis. Psoas major acts with iliacus as a powerful flexor of the thigh and trunk, while quadratus lumborum is a trunk extensor and a lateral flexor of the vertebral column and pelvis unilaterally (26). Cranially-oriented lumbar transverse processes with large attachment areas give all of these muscles effective leverage in acting on the vertebral column to assist the erector spinae in supporting the pelvis during upright posture and bipedalism (26, 45) and during ape-like vertical climbing (42). The lumbar vertebral morphology of *A. sediba*, therefore, is that of a biped equipped with powerful trunk musculature.

Williams et al.(18) predicted strong lumbar lordosis (“hyperlordosis”) in MH2 based on the sum wedging values of the penultimate and ultimate lumbar vertebrae. In contrast, Been et al. (21) estimated lumbar lordosis angle using pelvic incidence from a pelvis reconstruction(20) and found MH2 to produce the least lordotic lumbar column of the australopiths in their sample, falling well below modern human values and within the distribution of Neandertals. Neandertals are thought to be “hypolordotic,” or characterized by a relatively straight, non-lordotic lumbar column(21). Therefore, current interpretations of lumbar curvature of *A. sediba* range from hyperlordotic to hypolordotic. Here, we report that the pattern of vertebral wedging of MH2 is similar to *A. africanus* individuals at upper and middle lumbar levels, and more similar to *Homo* at lower lumbar levels (Fig. 6). Specifically, wedging values for MH2 fall between those of female *A. africanus* and modern human males at the penultimate lumbar level and produce the most dorsally-wedged last lumbar vertebra of known fossil hominins, approached only by the Kebara 2 Neandertal. Like Kebara 2, the strong dorsal (lordotic) wedging of MH2’s last lumbar vertebra is likely a reciprocate of strong ventral (kyphotic) wedging in the upper lumbar column (Fig. 6). However, MH2 demonstrates much less ventral wedging than Neandertals and produces a human-like overall sum of wedging angles, falling between modern human male and female means. Therefore, it seems likely that MH2 and some Neandertals demonstrate strong dorsal wedging at the last lumbar level for different reasons. It was suggested previously that the morphology of the MH2 last lumbar is part of a kinematic chain linked to hyperpronation of the foot(11, 18). With the absence of soft tissue contributions to the kinematic chain (i.e., intervertebral discs), formal biomechanical testing is beyond the scope of the current paper; however, our results suggest that MH2 was probably neither hypolordotic nor hyperlordotic and produces the most human-like sum of lumbar wedging angles of any early hominin yet known.

Modern humans are characterized by a pyramidal configuration of the lumbar inter-articular facet joints such that upper lumbar articular facet joints are transversely more narrowly spaced than those of lower lumbar vertebrae and especially compared to the lumbosacral inter-articular facet joints(44, 46). Together with vertebral body and intervertebral disc wedging, this progressive widening facilitates the adoption of lordotic posture during ontogeny and allows for the imbrication of the inferior articular facets of a superjacent vertebra onto the laminae of the subjacent vertebra during hyperextension of the lower back(18, 47). Like modern humans, known fossil hominin lumbar vertebrae bear “imbrication pockets,” mechanically induced fossae positioned just caudal to the superior articular facets on the lamina(18, 47), providing direct evidence for lumbar hyperextension and lordosis. Inadequate spacing of lower lumbar inter-articular facets in modern humans can result in spondylolysis, fracture of the *pars interarticularis* and potential separation of the affected vertebra’s spinous process and inferior articular processes(46). Lack of the progressive widening of inter-articular facets of lower lumbar vertebrae in our closest living relatives, the African apes, begs the question of when the pyramidal configuration evolved and to what extent various fossil hominins demonstrated this trait. Latimer and Ward(47) documented its presence in *Homo erectus*. Its presence has been demonstrated qualitatively in *A. afarensis* and *A. africanus*(26, 30), and its presence in *A. sediba* could be inferred previously based on the articulated last lumbar vertebrae and sacrum of MH2(18). Here we show that australopiths, including MH2, fall at the low end of modern human variation and differ from great apes in having appreciably wider inter-articular facets at the lumbosacral junction than higher in the lumbar column (Fig. 7).

Previous work has shown that the adult, presumed female individual from Malapa (MH2) demonstrates clear adaptations to bipedal locomotion (11–13, 18, 19), as do other australopiths, despite their retention of features linked to suspensory behavior and other arboreal proclivities (5–8, 10–17). The new fossils here reinforce these conclusions, signaling a lower back in MH2 as that of an upright biped equipped with powerful trunk musculature potentially used in both terrestrial and arboreal locomotion. The recovery and study of new fossil material, including juvenile material such that the ontogeny of bipedal features can be examined (48, 49), along with experimental biomechanical work and additional comparative analyses, will allow for testing hypotheses of form and function in the hominin fossil record.

## Materials and Methods

### 2D analyses (wedging angle, pyramidal configuration, lumbosacral joint)

Original fossil material was studied in all cases with the exception of the Neandertal fossils (Kebara 2 and Shanidar 3), for which high-quality were used. The remaining fossils were studied at the University of the Witwatersrand, the Ditsong National Museum of Natural History, the National Museum of Ethiopia, the National Museums of Kenya, Musée de l’Homme, and the Natural History Museum (London). Our comparative sample varied among 2D analyses but in total consisted of 70 chimpanzees (*Pan troglodytes*, 56 western gorillas, 17 orangutans (*Pongo* sp.), and 80 modern humans (*Homo sapiens*). To ensure that adequate space between elements was taken into account, we only included great apes with four lumbar vertebrae. Eastern gorillas (*Gorilla beringei*), which mostly possess just three lumbar vertebrae(50) are not included here, nor are other great ape individuals with only three lumbar vertebrae. The human sample includes data from an archaeological sample representing individuals from Africa, Asia-Pacific, and South America studied at the American Museum of Natural History, Musée de l’Homme, Musée Royal de l’Afrique Centrale, the Natural History Museum, and the University of the Witwatersrand. Measurements (listed in the Supplementary Note 1) were collected with Mitutoyo digital calipers and recorded at 0.01 mm; however, we report measurements at 0.1 mm.

Following DiGiovanni et al.(51), we use the wedging angle equation (Table 1) to calculate wedging angles for lumbar vertebrae 2-5. We also sum those values into a sum wedging value. Since the focus of this analysis is on fossil hominins, we only include well-sampled chimpanzees and western gorillas from the comparative sample, and we only report their sum wedging values. Confidence intervals (95%) were calculated for extant taxa using a Bootstrap resampling procedure with 9999 replicates.

Inter-articular facet spacing was measured across the lateral borders of the inferior articular facets of the L1 (in great apes) or L2 (in hominins) vertebra and on the last lumbar vertebra. Due to preservation, this measurement was estimated from the superior articular facets of the L3 vertebra and/or the sacrum in a selection of fossils (A.L. 288-1, Sts 14). In instances of partial preservation, the relevant adjacent elements were articulated to estimate the measurement (MH2, StW 431; KNM-WT 15000). An index was created by dividing the last lumbar-sacrum interarticular facet mediolateral width by that of the upper lumbar vertebrae.

### 3D reconstruction and geometric morphometric analysis

For 3D GM analyses, we used subsets of middle lumbar vertebrae (L3 in humans, L2 in great apes) that were scanned at the aforementioned institutions using an Artec Space Spider 3D scanner. Thirty-six modern humans, 28 chimpanzees, 26 western gorillas, and eight orangutans were included. For this analysis, we also utilized a sample of 23 bonobos (*Pan paniscus*) and seven eastern gorillas (*Gorilla beringei*).

U.W.88-233 is a complete lumbar vertebra, but it is partially encased in breccia, which obscures some morphologies. The entire new lumbar block (U.W.88-232-234) was mCT scanned at the University of the Witwatersrand. The high-resolution mCT scans were processed to yield virtual 3D models. Each vertebra was segmented using Amira 6.2. After importing mCT scan slices (TIFF files) and creating a volume stack file (.am), an *Edit New Label Field* module was attached to the stack file. Voxels were selected and assigned to each model separately using the *magic wand* and *brush* tools after verification in all three orthogonal views. A *Generate Surface* module was used to produce a *labels* file (.labels.am) once an individual element was completely selected. A 3D surface model was created from the *labels* file using an unconstrained smoothing setting of 5. Models of each element were then saved as polygon (.ply) files. Using GeoMagic Studio software, broken portions of U.W.88-233 were refit and the specimen was reconstructed accordingly. The affected portions of the neural arch were pulled dorsally to refit the fractured portion of the left lamina; additionally, the broken and deflected lumbar transverse process was refit. The result is a reconstructed 3D model (Supplementary Fig. 2).

Due to crushing of the right superior articular facet, we collected landmarks on the left side of U.W.88-233 and our comparative sample of middle lumbar vertebrae (Table 1). Our 3D landmark set consisted of 48 landmarks distributed across the vertebra to reflect the gross morphology (Supplementary Note 2). Landmarks were collected using the *Landmarks* tool in Amira on the surface model of U.W.88-233 and on 3D models of middle lumbar vertebrae produced using Artec Studio 14 software.

We used the geomorph package(52) in RStudio (the R Project for Statistical Computing) to carry out 3D GM analyses. Raw landmarks were imported into PAleontological STatistics (PAST) software and principal components analysis (PCA) was carried out to check for outliers. The geomorph package was then used to subject the raw landmark data to Generalized Procrustes analysis (GPA) to correct for position, rotation, and size adjustment. The GPA shape scores were then subjected to PCA using the covariance matrix. We evaluated the effects of centroid size on shape using Procrustes distance-based Analysis of Variance (ANOVA) on GPA shape scores as implemented in the geomorph package(23, 52).

We carried out two analyses: one on the full set of 48 landmarks in which only complete (reconstructed) fossils (U.W.88-233, Sts 14c, Shanidar 3) were included, and one on a 37 landmark subset with 11 landmarks on the transverse and spinous processes removed so that additional, less well-preserved fossils could be included (A.L. 288-1aa/ak/al, StW 431, Kebara 2).

## Acknowledgments

We thank the University of the Witwatersrand and the Evolutionary Studies Institute, as well as the South African National Centre of Excellence in PalaeoSciences and Bernhard Zipfel and Sifelani Jirah for curating the *A. sediba* material and allowing us access to it and to fossil comparative material in the Phillip V. Tobias Fossil Primate and Hominid Laboratory. We are grateful to Kudakwashe Jakata and Kris Carlson for assistance with microCT scanning. We thank the following individuals for curating and providing access to comparative materials in their care: Mirriam Tawane, Stephany Potze, and Lazarus Kgasi (Ditsong National Museum of Natural History); Brendon Billings and Anja Meyer (Dart Collection, University of the Witwatersrand); Yonas Yilma, Tomas Getachew, Jared Assefa, and Getachew Senishaw (National Museum of Ethiopia and Authority for Research and Conservation of Cultural Heritage); Emma Mbua (National Museums of Kenya); Véronique Laborde, Liliana Huet, Dominique Grimaud-Hervé, and Martin Friess (Musée de l’Homme); Rachel Ives (the Natural History Museum, London); Wim Wendelen and Emmanuel Gilissen (Musée Royal de l’Afrique Centrale); Lyman Jellema and Yohannes Haile-Selassie (Cleveland Museum of Natural History); and Gisselle Garcia, Ashley Hammond, Eileen Westwig, Eleanor Hoeger, Aja Marcato, Brian O’Toole, Marisa Surovy, Sarah Ketelsen, and Neil Duncan (American Museum of Natural History). Bill Kimbel and Chris Stringer facilitated access to fossils, and Erik Trinkaus shared high-quality casts of Kebara 2. We thank the South African Heritage Resource agency for the permits to work at Malapa and the Nash family for granting access to the site and continued support of research on their reserve, along with the South African Department of Science and Technology, the Gauteng Provincial Government, the Gauteng Department of Agriculture, Conservation and Environment and the Cradle of Humankind Management Authority, the South African National Research Foundation and the African Origins Platform, the National Geographic Society, the Palaeontological Scientific Trust (PAST), and the University of Witwatersrand’s Schools of Geosciences and Anatomical Sciences and the Bernard Price Institute for Paleontology for support and facilities, as well as our respective universities. We acknowledge the Leakey Foundation for funding support.

## Author contributions

S.A.W., T.C.P., M.R.M., T.K.N., C.Y., and D.G.M. contributed to data collection, S.A.W. and T.C.P. performed the analyses, and S.A.W., T.C.P., and R.D.V.M. wrote the manuscript, with contributions and feedback from all authors.

## SUPPLEMENTARY INFORMATION

### Supplementary Notes

#### Supplementary Note 1: Linear and angular measurements

The following measurements were taken on original fossil material or rendered surface models generated from high-resolution mCT scans (see descriptions in Results). Below,

1. Vertebral body superior transverse width (MαW7): defined in Bräuer (55) as the superior vertebral body transverse diameter at the most laterally projecting points.
2. Vertebral body superior dorsoventral length (Mα4): defined in Bräuer (55) as the superior vertebral body DV diameter measured at the sagittal midline.
3. Vertebral body inferior transverse width (Mα8): defined in Bräuer (55) as the inferior vertebral body transverse diameter at the most laterally projecting points.
4. Vertebral body inferior dorsoventral length (MαW5): defined in Bräuer (55) as the inferior vertebral body DV diameter measured at the sagittal midline.
5. Vertebral body superior-inferior ventral height (MαW1): defined in Bräuer (55) as the ventral SI height of the vertebral body at the sagittal midline.
6. Vertebral body superior-inferior dorsal height (MαW2): defined in Bräuer (55) as the dorsal SI height of the vertebral body at the sagittal midline.
7. Vertebral body wedging angle: calculation provided in DiGiovanni and colleagues (51) as
  1. *[arctangent ((((SI dorsal height-superior DV length)/2)/SI ventral height)*2)]*.
8. Spinal canal dorsoventral length (MαW10): defined in Bräuer (55) as DV spinal canal diameter measured at the sagittal midline.
9. Spinal canal transverse width (MαW11): defined in Bräuer (55) as transverse spinal canal diameter measured at the roots of the vertebral arch.
10. Superior-inferior inter-articular facet height: SI inter-articular facet distance, measured from the most superior aspect of the superior articular facet (SAF) to the most inferior aspect of the inferior articular facet (IAF) on the same side. The right side is measured unless it is broken or pathological, in which case the left side is measured. If the two sides are asymmetrical and one is not pathological, the mean is recorded.
11. Maximum inter-SAF transverse width: maximum (max.) inter-SAF distance, measured from the lateral aspect of one superior articular facet to the lateral aspect of the other.
12. Minimum inter-SAF transverse width: minimum (min.) inter-SAF distance, measured from the medial aspect of one superior articular facet to the medial aspect of the other.
13. Maximum inter-IAF transverse width: maximum (max.) inter-IAF distance, measured from the lateral aspect of one inferior articular facet to the lateral aspect of the other.
14. Minimum inter-IAF width: minimum (min.) inter-IAF distance, measured from the medial aspect of one inferior articular facet to the medial aspect of the other.
15. SAF superior-inferior height: SAF SI diameter, measured at the sagittal midline. The right side is measured unless it is broken or pathological, in which case the left side is measured. If the two sides are asymmetrical and one is not pathological, the mean is recorded.
16. SAF transverse width: SAF transverse diameter, measured from the most medial to the most lateral border of the articular surface. The right side is measured unless it is broken or pathological, in which case the left side is measured. If the two sides are asymmetrical and one is not pathological, the mean is recorded.
17. IAF superior-inferior height: IAF SI diameter, measured at the sagittal midline. The right side is measured unless it is broken or pathological, in which case the left side is measured. If the two sides are asymmetrical and one is not pathological, the mean is recorded.
18. IAF transverse width: IAF transverse diameter, measured from the most medial aspect to the most lateral border of the articular surface. The right side is measured unless it is broken or pathological, in which case the left side is measured. If the two sides are asymmetrical and one is not pathological, the mean is recorded.
19. Spinous process angle (MαW12): defined in Bräuer (55) as the angle that is formed from the superior surface of the vertebral body and the upper edge of the spinous process. We modify this measurement slightly by measuring the angle along its long axis, which allows for the inclusion of fossils with a damaged or missing superior edge of the spinous process. An angle of 180° is equivalent to a spinous process with a long axis parallel to the superior surface of the vertebral body (i.e., horizontal or neutral in orientation).
20. Spinous process length (MαW13): defined in Bräuer (55) as the distance from the top edge of the vertebral arch to the most dorsal tip of the spinous process.
21. Spinous process terminal transverse width: transverse breadth of the dorsal tip of the spinous process, measured at its maximum dimension.
22. Spinous process terminal superior-inferior height: SI diameter of the dorsal tip of the spinous process, measured at the mediolateral midline of the spinous process.
23. Transverse process superior-inferior base height: SI diameter of the transverse process at its medial origin from the pedicle and/or vertebral body.
24. Transverse process dorsoventral angle: the dorsoventral angle that is formed from the sagittal midplane of the vertebra to the long axis of the transverse process, along the middle from its base to its tip. This contrasts with Ward et al. (56), where the transverse process angle is measured along the costal facet. We justify the use of the long axis because lower thoracic vertebrae and lumbar vertebrae lack costal facets. The right side is measured unless it is broken or pathological, in which case the left side is measured. If the two sides are asymmetrical and one is not pathological, the mean is recorded.
25. Transverse process length: the distance from the internal edge of the spinal canal at its closest point to the base of the transverse process to the tip of the transverse process. The right side is measured unless it is broken or pathological, in which case the left side is measured. If the two sides are asymmetrical and one is not pathological, the mean is recorded.
26. SAF orientation: the angle formed between the sagittal midplane of the vertebra to the medial and lateral edges of the SAF. The right side is measured unless it is broken or pathological, in which case the left side is measured. If the two sides are asymmetrical and one is not pathological, the mean is recorded.
27. Pedicle superior-inferior height: SI diameter of the pedicle, measured at its midpoint. The right side is measured unless it is broken or pathological, in which case the left side is measured. If the two sides are asymmetrical and one is not pathological, the mean is recorded.
28. Pedicle transverse width: transverse breadth of the pedicle, measured at its midpoint. The right side is measured unless it is broken or pathological, in which case the left side is measured. If the two sides are asymmetrical and one is not pathological, the mean is recorded.
29. Pedicle dorsoventral length: DV diameter of the pedicle, measured anterior from its junction with the superior articular process to its junction with the dorsal edge of the vertebral body. The right side is measured unless it is broken or pathological, in which case the left side is measured. If the two sides are asymmetrical and one is not pathological, the mean is recorded.
30. Lamina superior-inferior height: SI dimension of the lamina, measured on one side between the spinous process and the SAF and the IAF. The right side is measured unless it is broken or pathological, in which case the left side is measured. If the two sides are asymmetrical and one is not pathological, the mean is recorded.
31. Lamina transverse width: transverse dimension of the lamina, measured at its minimum breadth across the *pars interarticularis*.

#### Supplementary Note 2: List of 3D landmarks

The following 3D landmarks were collected on middle lumbar vertebrae using AMIRA:

1. Superior vertebral body – ventral transverse midline
2. Superior vertebral body – central transverse midline
3. Superior vertebral body – dorsal transverse midline
4. Superior vertebral body – lateral sagittal midline
5. Superior vertebral body – ventro-lateral point
6. Superior vertebral body – dorso-lateral point (at ventral pedicle base)
7. Pedicle – superior midpoint
8. Pedicle – superior dorsal point (at ventral base of prezygapophysis)
9. Pedicle – medial midpoint
10. Pedicle – lateral midpoint
11. Pedicle – inferior midpoint
12. *Pars interarticularis* – dorsal midpoint
13. *Pars interarticularis* – ventral midpoint
14. Inferior vertebral body – ventral transverse midline
15. Inferior vertebral body – central transverse midline
16. Inferior vertebral body – dorsal transverse midline
17. Inferior vertebral body – lateral sagittal midline
18. Inferior vertebral body – ventro-lateral point
19. Inferior vertebral body – dorso-lateral point
20. Lumbar transverse process (LTP) – superior medial point (based of LTP at pedicle)
21. Lumbar transverse process – inferior medial point (based of LTP at pedicle)
22. Lumbar transverse process – superior mediolateral midpoint
23. Lumbar transverse process – inferior mediolateral midpoint
24. Lumbar transverse process – superior lateral point
25. Lumbar transverse process – inferior lateral point
26. Superior articular facet – cranial-most point
27. Mammillary process – dorsal-most extension
28. Superior articular facet – midpoint
29. Superior articular facet – caudal-most point
30. Spinous process – superior ventral point (at spinous process base)
31. Spinous process – superior sagittal midpoint
32. Spinous process – superior dorsal point (tip of spinous process)
33. Spinous process – inferior dorsal point (tip of spinous process)
34. Spinous process – inferior sagittal midpoint
35. Spinous process – inferior ventral point (at spinous process base)
36. Lamina – inferior midpoint
37. Inferior articular facet – cranial-most point
38. Inferior articular facet – midpoint
39. Inferior articular facet – caudal-most point
40. Inferior articular facet – medial-most point
41. Inferior articular facet – lateral-most point
42. Superior articular facet – medial-most point
43. Superior articular facet – lateral-most point
44. Vertebral body – ventral midpoint
45. Vertebral body – lateral midpoint
46. Lumbar transverse process – ventral extension of base at its midpoint
47. Spinous process – lateral-most extension of the spinous process tip
48. Lamina – superior midpoint

## Supplementary Figures

**Supplementary Figure 1.**
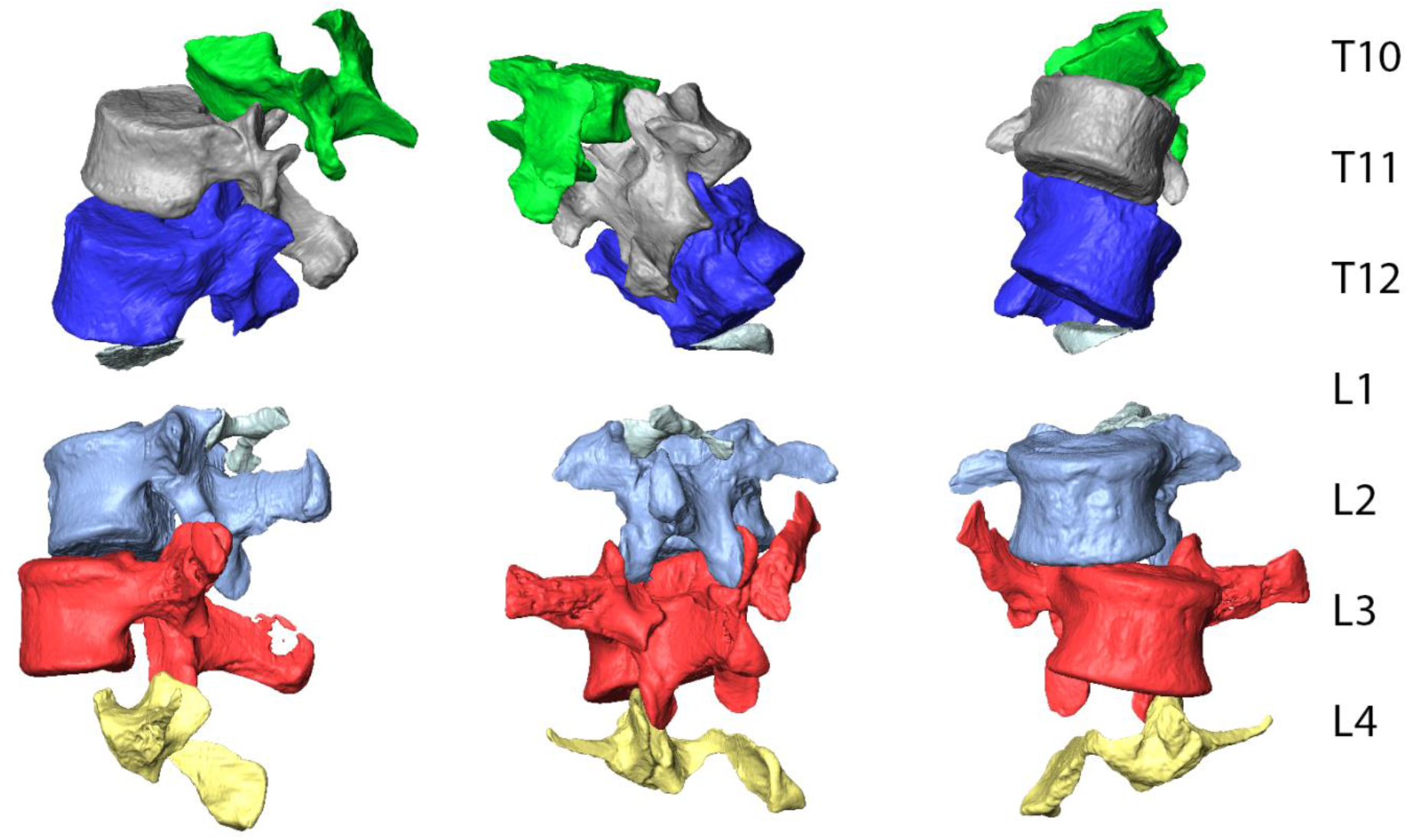
Surface renderings of lower thoracic and lumbar vertebrae generated from microCT scans. Two matrix blocks were microCT scanned: a lower thoracic block containing the antepenultimate (T10; U.W. 88-114), penultimate (T11; U.W. 88-43), last thoracic vertebra (T12; U.W. 88-44), and vertebral body fragment of the first lumbar vertebra (L1; U.W. 88-280), and the new lumbar block, containing the partial inferior articular facets of L1 (U.W. 88-281), second lumbar vertebra (L2; U.W. 88-232), third lumbar vertebra (L3; U.W. 88-233), and portions of the upper neural arch of the fourth lumbar vertebra (L4; U.W. 88-234). High quality surface models of these vertebrae can be downloaded on MorphoSource.org.

**Supplementary Figure 2.**
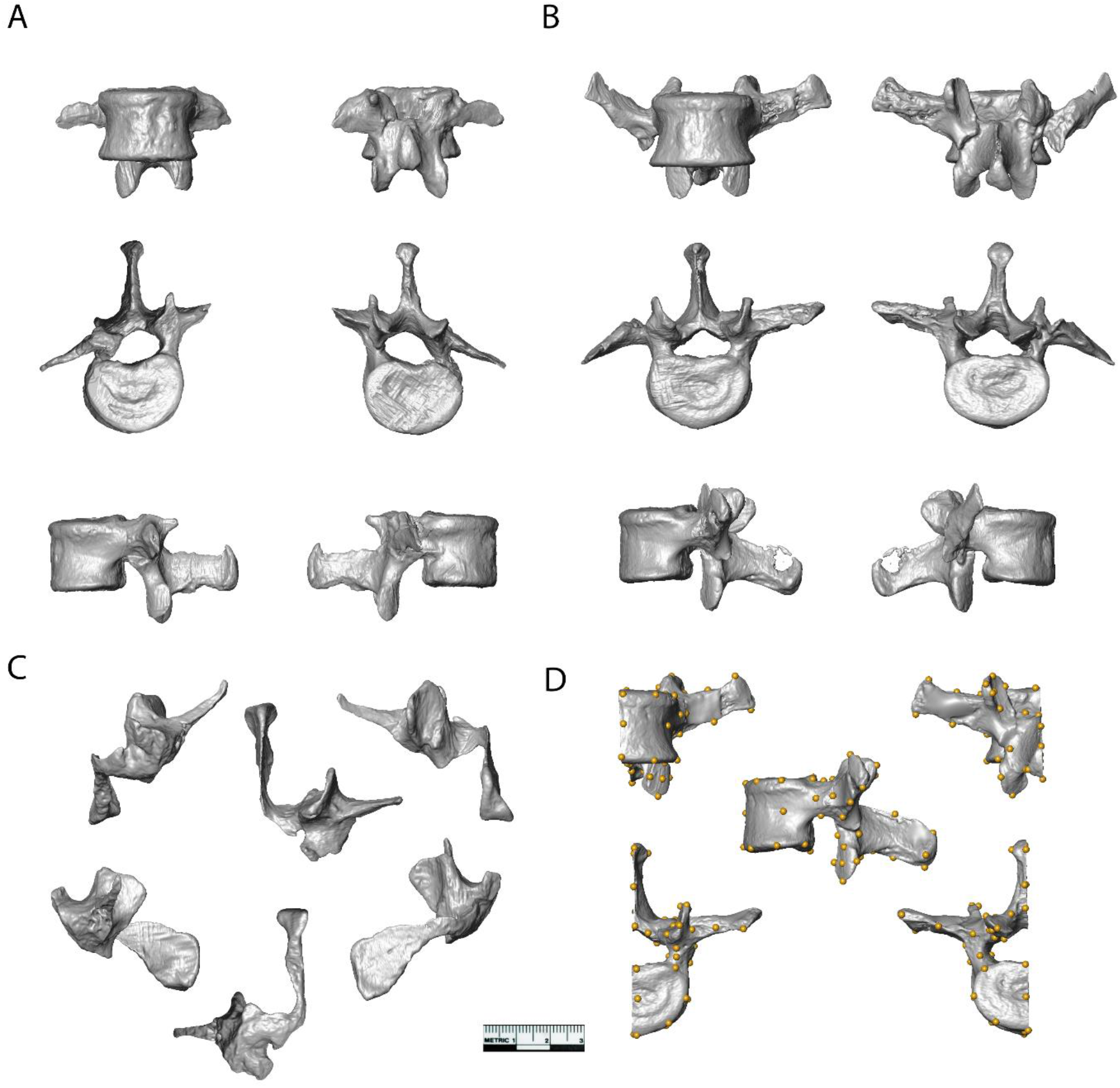
3D surface models of vertebrae from the new lumbar block. **(A)** U.W. 88-232 (L2) and **(B)** U.W. 88-233 (L3) shown in ventral (top left), dorsal (top right), superior (middle left), inferior (middle right), left lateral (bottom left), and right lateral (bottom right) views. **(C)** U.W. 88-234 (L4) in ventral (top left), dorsal (top right), superior (top middle), left lateral (bottom left), right lateral (bottom right), and inferior (bottom middle) views. **(D)** Left half of U.W. 88-233 showing the 48 landmarks used in 3D GM analyses.

**Supplementary Figure 3.**
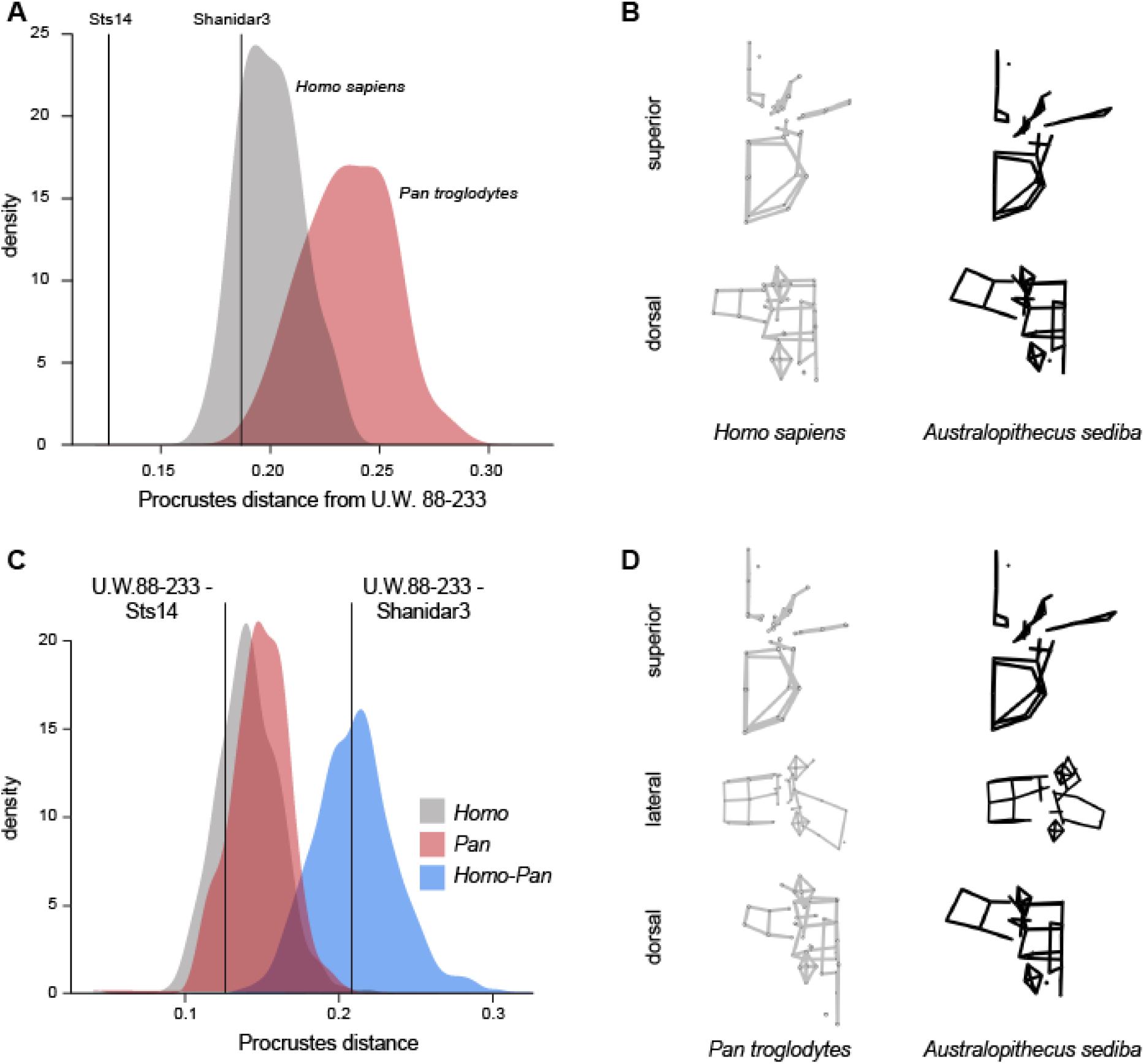
Procrustes distances and mean differences from U.W. 88-233. **(A)** Procrustes distances between U.W. 88-233 and extant and fossil middle lumbar vertebrae. Sts14 is closest to U.W. 88-233 in overall Procrustes distance. **(B)** Average modern human middle lumbar morphology compared to U.W. 88-233. **(C)** Procrustes distances within (*Homo, Pan*) and between (*Pan* and *Homo*) extant taxa and between U.W. 88-233 and other fossil hominin middle lumbar vertebrae. **(D)** Average *Pan* middle lumbar vertebra morphology compared to U.W. 88-233.

**Supplementary Figure 4.**
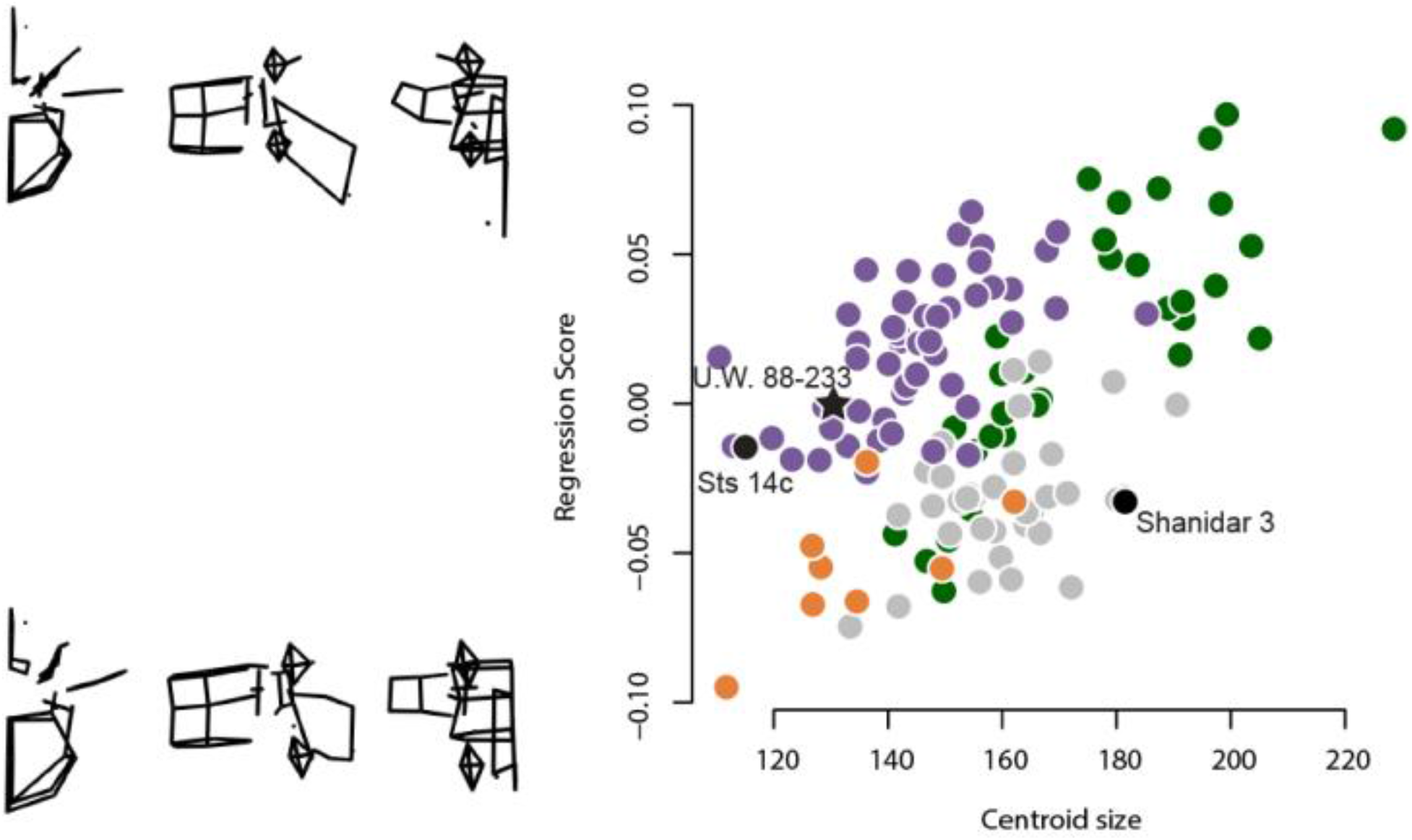
The effect of middle lumbar centroid size on shape. Wireframes represent the maximum and minimum shape predictions derived from a multivariate regression of shape coordinates on centroid size. Larger centroid sizes are associated with longer, more projecting spinous processes and transverse processes. The elongated and cranially oriented transverse processes of U.W. 88-233 (*A. sediba*) and Sts 14c (*A. africanus*) are not explained by centroid size. *Homo* = grey circles, *Pan* = purple circles, *Gorilla* = green circles, *Pongo* = orange circles.

## References

1. T. D. White, C. O. Lovejoy, B. Asfaw, J. P. Carlson, G. Suwa, Neither chimpanzee nor human, Ardipithecus reveals the surprising ancestry of both. Proceedings of the National Academy of Sciences USA 112, 4877–4884 (2015).

2. T. C. Prang, The African ape-like foot of Ardipithecus ramidus and its implications for the origin of bipedalism. eLife 8, e44433 (2019).

3. L. R. Berger et al., Australopithecus sediba: a new species of Homo-like australopith from South Africa. Science 328, 195–204 (2010).

4. P. Schmid et al., Mosaic morphology in the thorax of Australopithecus sediba. Science 340, 1234598 (2013).

5. T. C. Prang, Calcaneal robusticity in Plio-Pleistocene hominins: Implications for locomotor diversity and phylogeny. Journal of Human Evolution 80, 135–146 (2015).

6. T. C. Prang, Rearfoot posture of Australopithecus sediba and the evolution of the hominin longitudinal arch. Scientific Reports 5, 17677 (2015).

7. T. C. Prang, The subtalar joint complex of Australopithecus sediba. Journal of Human Evolution 90, 105–119 (2016).

8. T. W. Holliday et al., Body Size and Proportions of Australopithecus sediba. PaleoAnthropology 406-422, 422 (2018).

9. C. J. Dunmore et al., The position of Australopithecus sediba within fossil hominin hand use diversity. Nature Ecology & Evolution, 1–8 (2020).

10. T. R. Rein, T. Harrison, K. J. Carlson, K. Harvati, Adaptation to suspensory locomotion in Australopithecus sediba. Journal of Human Evolution 104, 1–12 (2017).

11. J. M. DeSilva et al., The lower limb and mechanics of walking in Australopithecus sediba. Science 340, 1232999 (2013).

12. B. Zipfel et al., The foot and ankle of Australopithecus sediba. Science 333, 1417–1420 (2011).

13. J. M. DeSilva et al., The Anatomy of the Lower Limb Skeleton of Australopithecus sediba. PaleoAnthropology 357, 405 (2018).

14. S. E. Churchill et al., The upper limb of Australopithecus sediba. Science 340, 1233477 (2013).

15. S. E. Churchill et al., The Shoulder, Arm, and Forearm of Australopithecus sediba. PaleoAnthropology 234, 281 (2018).

16. M. R. Meyer, S. A. Williams, P. Schmid, S. E. Churchill, L. R. Berger, The cervical spine of Australopithecus sediba. Journal of Human Evolution 104, 32–49 (2017).

17. A. G. Henry et al., The diet of Australopithecus sediba. Nature 487, 90–93 (2012).

18. S. A. Williams et al., The vertebral column of Australopithecus sediba. Science 340, 1232996 (2013).

19. S. A. Williams et al., The Vertebrae, Ribs, and Sternum of Australopithecus sediba. PaleoAnthropology 2018, 156–233 (2018).

20. J. M. Kibii et al., A partial pelvis of Australopithecus sediba. Science 333, 1407–1411 (2011).

21. E. Been, A. Gómez-Olivencia, P. A. Kramer, Lumbar lordosis in extinct hominins: implications of the pelvic incidence. American Journal of Physical Anthropology 154, 307–314 (2014).

22. P. H. Dirks et al., Geological setting and age of Australopithecus sediba from southern Africa. Science 328, 205–208 (2010).

23. C. Goodall, Procrustes methods in the statistical analysis of shape. Journal of the Royal Statistical Society B 53, 285–321 (1991).

24. A. G. Drake, C. P. Klingenberg, The pace of morphological change: historical transformation of skull shape in St Bernard dogs. Proceedings of the Royal Society B 275, 71–76 (2008).

25. P. Davis, Human lower lumbar vertebrae: some mechanical and osteological considerations. Journal of Anatomy 95, 337 (1961).

26. J. T. Robinson, Early Hominid Posture and Locomotion (University of Chicago Press, Chicago, 1972).

27. G. Pal, Weight transmission through the sacrum in man. Journal of Anatomy 162, 9–17 (1989).

28. B. Latimer, Ward, C. V., “The thoracic and lumbar vertebrae.” in 1993, A. Walker, Leakey, R., Ed. (Harvard University Press, Cambridge, MA, 1993), pp. 266-293.

29. L. Shapiro, Evaluation of “unique” aspects of human verterbal bodies and pedicles with a consideration of Australopithecus africanus. Journal of Human Evolution 25, 433–470 (1993).

30. C. O. Lovejoy, The natural history of human gait and posture: Part 1. Spine and pelvis. Gait & Posture 21, 95–112 (2005).

31. K. K. Whitcome, L. J. Shapiro, D. E. Lieberman, Fetal load and the evolution of lumbar lordosis in bipedal hominins. Nature 450, 1075–1078 (2007).

32. C. Fornai, V. Krenn, P. Mitteröcker, N. Webb, M. Haeusler, Sacrum morphology supports taxonomic heterogeneity of Australopithecus africanus at Sterkfontein Member 4. Research Square preprint doi:10.21203/rs.3.rs-72859/v 1, 1–18 (2020).

33. G. A. Macho et al., The partial skeleton StW 431 from Sterkfontein-Is it time to rethink the Plio-Pleistocene hominin diversity in South Africa? Journal of Anthropological Sciences 98, 73–88 (2020).

34. D. C. Johanson et al., Morphology of the Pliocene partial hominid skeleton (AL 288-1) from the Hadar formation, Ethiopia. American Journal of Physical Anthropology 57, 403–451 (1982).

35. M. R. Meyer, S. A. Williams, M. P. Smith, G. J. Sawyer, Lucy’s back: Reassessment of fossils associated with the A.L. 288-1 vertebral column. Journal of Human Evolution 85, 174–180 (2015).

36. D. C. Cook, J. E. Buikstra, C. J. DeRousseau, D. C. Johanson, Vertebral pathology in the Afar australopithecines. American Journal of Physical Anthropology 60, 83–101 (1983).

37. E. J. Slijper, Comparative biologic-anatomical investigations on the vertebral column and spinal musculature of mammals. Verh. Kon. Ned. Akad. Wet. (Tweede Sectie), 1–128 (1946).

38. L. Shapiro, “Functional morphology of the vertebral column in primates.” in Postcranial Adaptation in Nonhuman Primates, D. L. Gebo, Ed. (Northern Illinois University Press, Dekalb, IL, 1993), pp. 121–149.

39. W. J. Sanders, B. E. Bodenbender, Morphometric analysis of lumbar vertebra UMP 67-28: implications for spinal function and phylogeny of the Miocene Moroto hominoid. Journal of Human Evolution 26, 203–237 (1994).

40. M. C. Granatosky, C. E. Miller, D. M. Boyer, D. Schmitt, Lumbar vertebral morphology of flying, gliding, and suspensory mammals: implications for the locomotor behavior of the subfossil lemurs Palaeopropithecus and Babakotia. Journal of Human Evolution 75, 40–52 (2014).

41. S. A. Williams, G. A. Russo, Evolution of the hominoid vertebral column: the long and the short of it. Evolutionary Anthropology: Issues, News, and Reviews 24, 15–32 (2015).

42. P. Schmid, “The trunk of the australopithecines” in Origine (s) de la Bipédie chez les Hominidés, Senut, Y. Coppens, Eds. (Editions du CNRS, Paris, 1991), pp. 225–234.

43. L. Shapiro, Evaluation of” unique” aspects of human vertebral bodies and pedicles with a consideration of Australopithecus africanus. Journal of Human Evolution 25, 433–470 (1993).

44. W. J. Sanders, Comparative morphometric study of the australopithecine vertebral series Stw-H8/H41. Journal of Human Evolution 34, 249–302 (1998).

45. R. Waters, J. Morris, Electrical activity of muscles of the trunk during walking. Journal of Anatomy 111, 191–199 (1972).

46. C. V. Ward, B. Latimer, Human evolution and the development of spondylolysis. Spine 30, 1808–1814 (2005).

47. B. Latimer, C. V. Ward, “The thoracic and lumbar vertebrae” in The Nariokotome Homo erectus Skeleton, A. Walker, R. Leakey, Eds. (Harvard University Press, Cambridge, MA, 1993), pp. 266–293.

48. C. V. Ward, T. K. Nalley, F. Spoor, P. Tafforeau, Z. Alemseged, Thoracic vertebral count and thoracolumbar transition in Australopithecus afarensis. Proceedings of the National Academy of Sciences 114, 6000–6004 (2017).

49. T. K. Nalley, J. E. Scott, C. V. Ward, Z. Alemseged, Comparative morphology and ontogeny of the thoracolumbar transition in great apes, humans, and fossil hominins. Journal of Human Evolution 134, 102632 (2019).

50. S. A. Williams, A. Gómez-Olivencia, D. R. Pilbeam, “Numbers of vertebrae in hominoid evolution” in Spinal Evolution, E. Been, A. Gómez-Olivencia, P. A. Kramer, Eds. (Springer, 2019), pp. 97–124.

51. B. F. DiGiovanni, P. V. Scoles, B. M. Latimer, Anterior extension of the thoracic vertebral bodies in Scheuermann’s kyphosis: an anatomic study. Spine 14, 712–716 (1989).

52. D. C. Adams, E. Otárola-Castillo, geomorph: an R package for the collection and analysis of geometric morphometric shape data. Methods in Ecology and Evolution 4, 393–399 (2013).

53. T. R. Pickering, J. L. Heaton, R. Clarke, D. Stratford, Hominin vertebrae and upper limb bone fossils from Sterkfontein Caves, South Africa (1998–2003 excavations). American Journal of Physical Anthropology 168, 459–480 (2019).

54. J. Hawks et al., New fossil remains of Homo naledi from the Lesedi Chamber, South Africa. Elife 6, e24232 (2017).

## Supplementary References

12. B. Zipfel et al., The foot and ankle of Australopithecus sediba. Science 333, 1417–1420 (2011).

14. S. E. Churchill et al., The upper limb of Australopithecus sediba. Science 340, 1233477 (2013).

17. A. G. Henry et al., The diet of Australopithecus sediba. Nature 487, 90–93 (2012).

20. J. M. Kibii et al., A partial pelvis of Australopithecus sediba. Science 333, 1407–1411 (2011).

27. G. Pal, Weight transmission through the sacrum in man. Journal of Anatomy 162, 9–17 (1989).

42. P. Schmid, “The trunk of the australopithecines” in Origine (s) de la Bipédie chez les Hominidés, B. Senut, Y. Coppens, Eds. (Editions du CNRS, Paris, 1991), pp. 225–234.

55. G. Bräuer (1988) Osteometrie. In (R. Knussmann, Ed.) Anthropologie. Handbuch der vergleichenden Biologie des Menschen. (Stuttgart: Gustav Fischer Verlag).

56. C. V. Ward, W. H. Kimbel, E. H. Harmon, D. C. Johanson, New postcranial fossils of Australopithecus afarensis from Hadar, Ethiopia (1990–2007). Journal of Human Evolution 63, 1–51 (2012).

